# *PCDHGC3* silencing promotes clear cell renal cell carcinoma metastasis via mTOR/HIF2α and lipid metabolism reprogramming

**DOI:** 10.1101/2024.08.26.609687

**Authors:** Lucía Celada, Tamara Cubiella, Laura Salerno, Jaime San-Juan-Guardado, Eduardo Murias, Marina Da Silva Torres, Álvaro Suárez-Priede, Joshua A. Weiner, Helena Herrada-Manchón, M. Alejando Fernández, María-Dolores Chiara

## Abstract

Clustered protocadherins (cPCDH) are widely expressed in the nervous system with known functions, but their roles in cancer, particularly metastasis, are largely unexplored. Our previous research revealed that epigenetic silencing of *PCDHGC3* is linked to decreased survival in neuroendocrine cancer patients. This study investigates *PCDHGC3*’s role in clear cell renal cell carcinoma (ccRCC). We found that decreased *PCDHGC3* expression is associated with lower survival and advanced disease stage in ccRCC patients. shRNA-mediated *PCDHGC3* silencing in renal cancer cell lines significantly increased cell proliferation, invasion, and survival. In orthotopic mouse models, *PCDHGC3* silencing promoted metastasis. The mTOR and HIF2α pathways were identified as downstream targets activated by *PCDHGC3* loss. Inhibition of these pathways counteracted the effects of *PCDHGC3* silencing, highlighting their importance in tumor progression. Proteomic and metabolomic analyses showed that *PCDHGC3* silencing led to overexpression of proteins involved in fatty acid and cholesterol synthesis, increasing lipid droplets and shifting lipid metabolism. This metabolic reprogramming characterizes aggressive ccRCC. Our study emphasizes *PCDHGC3*’s impact on ccRCC metastasis and suggests mTOR or HIF2α inhibitors as potential therapies for *PCDHGC3*-deficient patients.

## Introduction

Renal cell carcinoma (RCC) is the predominant form of kidney cancer in adults, encompassing about 95% of reported cases (Hsieh *et al*, 2017). This cancer holds a significant global prevalence standing among the top 10 most common cancers. Clear cell renal cell carcinoma (ccRCC) represents the most prevalent subtype, comprising around 75% of cases. This cancer is distinguished by the characteristic clear appearance of its cells when observed under a microscope, a feature attributed to the accumulation of cholesterol and other neutral lipids (Qi *et al*, 2021). Despite surgical removal remaining the primary curative approach for localized ccRCC, the challenge lies in addressing the complexities associated with recurrent and metastatic cases (Angulo & Shapiro, 2019). Advances in understanding the genetic underpinnings have paved the way for targeted therapies.

The most prevalent genetic disorder in patients with ccRCC affects the von Hippel-Lindau (VHL) gene(Kaelin, 2007). Inactivation of VHL precludes ubiquitination and degradation of hypoxia-inducible factors (HIFs), leading to accumulation of HIF and the subsequent upregulation of vascular endothelial growth factor (VEGF) and other protumorigenic pathways (Cockman *et al*, 2000; Maxwell *et al*, 1999). In addition to the VHL-HIF pathway, metabolic reprogramming is induced by the activation of the PI3K-AKT-mTOR (mammalian target of rapamycin) pathway (Hager *et al*, 2011). Given this understanding, current treatment options for patients with advanced and metastatic ccRCC are limited to targeted therapies that inhibit VEGF receptors, mTOR, and immunotherapy with checkpoint inhibitors (Meng *et al*, 2023; Wilson *et al*, 2024). More recently, small molecules targeting HIF2α have become accepted drugs for advanced ccRCC (Nguyen *et al*, 2024),(Expanding ‘Practice-Changing’ Belzutifan’s Reach in RCC, 2023).

Metastasis development entails the epithelial-to-mesenchymal transition of cancer cells, promoting the emergence of highly mobile and invasive mesenchymal cells compared to their epithelial counterparts. Consequently, in ccRCC, it is common to observe cancer cells that have lost expression of the epithelial cell biomarker, E-cadherin, a crucial component in cell-to-cell adhesion (Jang *et al*, 2021). This loss facilitates the emergence of motile, free cells capable of invading other target organs. Additionally, epigenetic silencing of the clustered protocadherin (c*PCDH*) genes have been identified in Wilms’ tumor, a pediatric kidney tumor (Dallosso *et al*, 2009) and colorectal cancer(Dallosso *et al*, 2012a). While *cPCDH* genes are extensively expressed in the nervous system, where they have been predominantly studied (Peek *et al*, 2017; Flaherty & Maniatis, 2020; Mountoufaris *et al*, 2018; Keeler *et al*, 2015a) , their role in other tissues remains poorly understood in terms of physiology and pathology. Recent findings have illuminated the potential of *cPCDH* as tumor suppressors (Vega-Benedetti *et al*, 2019; Dallosso *et al*, 2012a). Notably, within these genes, the methylation-dependent silencing of *PCDHGC3* has been shown to exert a significant impact on the aggressive behavior of some cancer types, including paraganglioma, pheochromocytoma (Bernardo-Castiñeira *et al*, 2019), colorectal cancer (Dallosso *et al*, 2012a), and gastrointestinal neuroendocrine carcinoma (Cubiella *et al*, 2024).

In this study, we examined the role of *PCDHGC3* in metastatic ccRCC. Our investigation reveals that the knockdown of this gene exerts a profound influence on the growth and metastatic behavior of ccRCC-derived cells and tumors, unraveling a significant mechanism involving the activation of the mTOR and HIF2α pathways and lipid metabolism reprogramming. This exploration sheds light on the molecular pathways underlying the metastatic cascade in ccRCC, providing valuable insights for potential therapeutic interventions.

## Results

### Distinct gene expression control of *PCDHGC3* among clustered *PCDH* genes in ccRCC

Analysis of mRNA sequencing data in ccRCC samples from the TCGA database revealed active transcription of all *cPCDH* genes in this type of tumor, with *PCDHGC3* being the most highly expressed (Fig 1A). Subsequent analysis revealed that decreased expression of *PCDHGC3* showed a strong association with advanced disease (stage IV) (p=0.0059) and low overall survival rates (p<0.0001) (Fig 1B and C). This association was not observed with *PCDHGC4,* the downstream gene within the C-type *PCDHG* cluster (Fig EV1).

**Figure 1.**
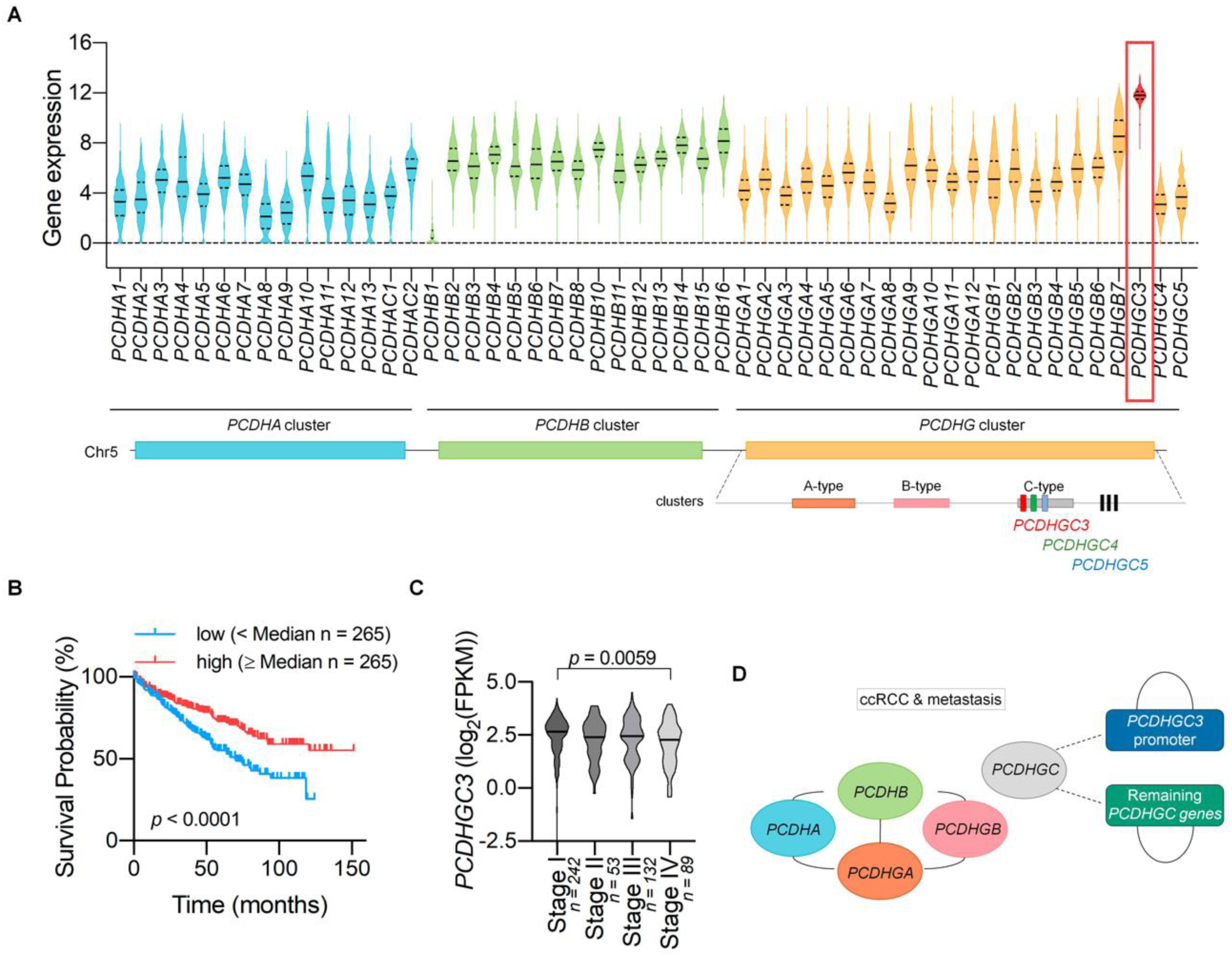
*PCDHGC3* expression in ccRCC and its correlation with disease outcome. (A) Expression levels of *cPCDHA*, *cPCDHB* and *cPCDHG* genes in ccRCC samples included in the TCGA database. An illustration in the bottom depicts the three *PCDH* gene clusters located at chromosome 5. The genomic organization of the *PCDHGC* cluster is depicted, consisting of 5′-proximal variable exons (represented as rectangles) specifically encoding *PCDHGC3* (in red), *PCDHGC4* (in green), or *PCDHGC5* (in blue), which splice to three exons (depicted as black rectangles) shared among the *PCDHG* genes. (B) Kaplan–Meier curves for ccRCC patients according to the *PCDHGC3* mRNA levels. (C) Association between *PCDHGC3* expression and disease stage. (D) Associations between CpG methylation patterns of *cPCDHG* genes in metastatic ccRCCs. Analysis was performed with the Autodiscovery software.

Each *cPCDH* gene has its own promoter and is known to be regulated by DNA methylation mechanisms. Our previous studies highlighted the significance of *PCDHGC3* methylation in metastatic paraganglioma (Bernardo-Castiñeira *et al*, 2019) and gastrointestinal neuroendocrine carcinoma (Cubiella *et al*, 2024) correlating with low survival rates. In our investigation of DNA methylation levels across *cPCDH* genes in ccRCC tumors compared to non-tumoral samples, we observed a notable elevation in methylation levels throughout the entire cluster (Fig EV1). In metastatic ccRCC, we observed coordinated methylation patterns among gene promoters within the *PCDHA*, *PCDHB*, and A and B-type *PCDHG* clusters. However, the methylation pattern of C-type *PCDHG* genes stood out distinctly, showing no correlation with that of other *cPCDH* promoters (Fig 1D). In fact, detailed analysis of the *PCDHG* cluster revealed significant hypermethylation levels in promoter regions of A-type and B-type genes in tumor versus normal tissues whereas very subtle hypermethylation levels were detected in the *PCDHGC3* gene promoter. In addition, we observed a strong inverse correlation between the expression levels of A-type and B-type genes within the *PCDHG* cluster and their corresponding promoter methylation levels (Fig EV1) while this correlation was less evident with *PCDHGC3* gene.

Taken together, these findings underscore the distinctive and clinically significant role of *PCDHGC3* in ccRCC, shedding light on the potential clinical significance of its specific regulation in this disease.

### *PCDHGC3* knockdown increases ccRCC growth

To understand the role of *PCDHGC3* role in ccRCC, we knocked down *PCDHGC3* expression in 786-O and RCC4 ccRCC-derived cell lines (hereafter termed C3KD) using three sequence-independent short hairpin RNAs (shRNAs). Cells expressing nonspecific shRNA were used as negative controls (CT cells). C3KD cell lines expressed significantly lower levels of *PCDHGC3* than CT cells (Fig 2A). The ability of these cells lines to grow was then evaluated using an MTS cell proliferation assay. We found a 1.27-fold and 1.25-fold increase in proliferation for C3KD-786-O and C3KD-RCC4 cells, respectively, compared to their respective controls (Fig 2B) corroborating our previous results (Bernardo-Castiñeira *et al*, 2019).

**Figure 2.**
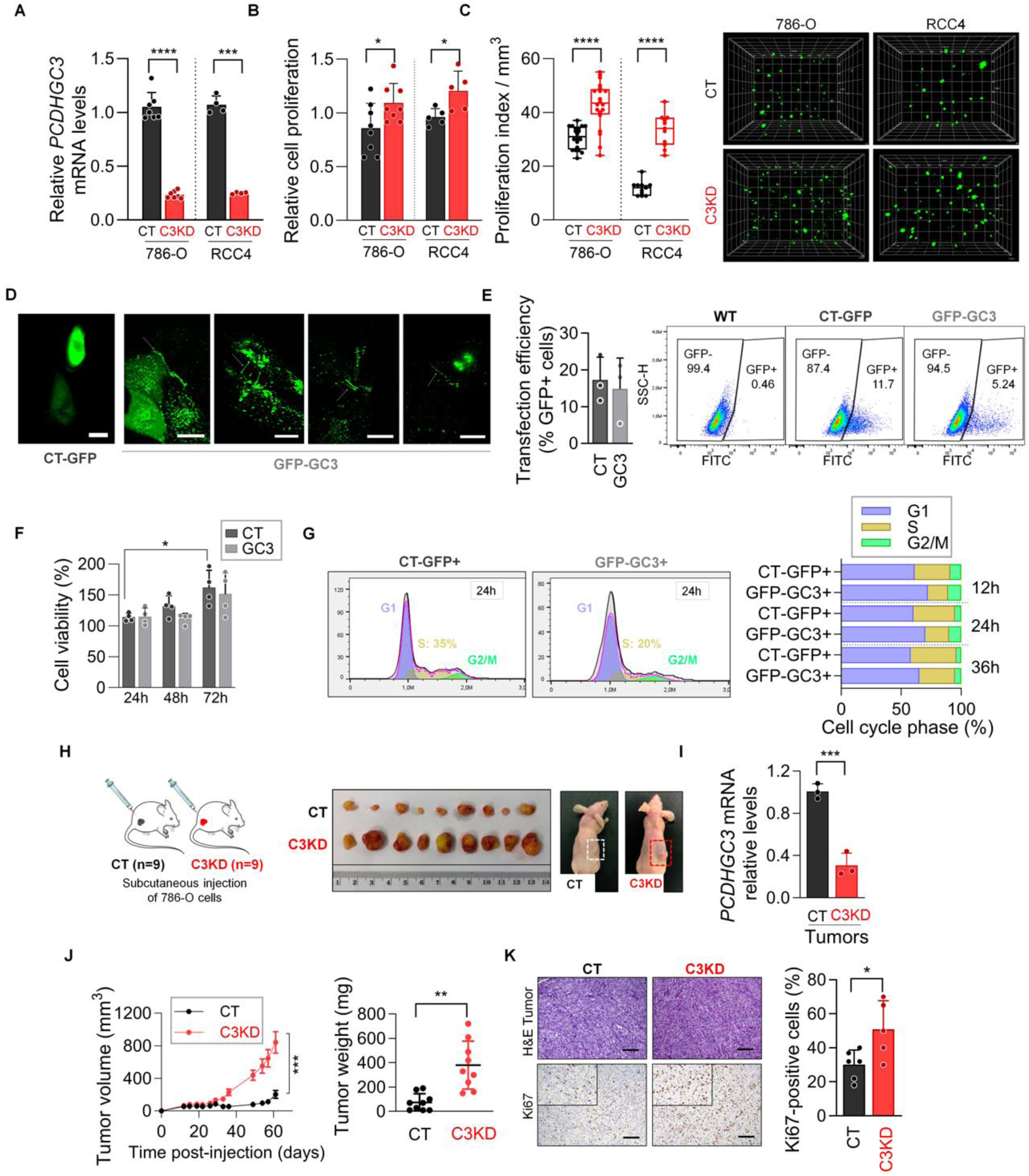
*PCDHGC3* knockdown promotes renal cancer cell proliferation in vitro and in vivo. (A) Relative *PCDHGC3* mRNA levels determined by RT-qPCR in 786-O and RCC4 cells upon shRNA-*PCDHGC3* transfection. (B) Cell proliferation of the indicated CT and C3KD cells assessed by MTS assay over a 72-h period. (C) Proliferation index of the indicated CT and C3KD following printing and incubation for a 14-day period. Cell counting was performed on Z-stack images (15-20 sections with an interval of 20 μm between them) at randomly chosen positions within the scaffold. Representative fluorescence XYZ projections of CT or C3KD printed cells at day 14 after printing are shown on the right. (D) Fluorescence microscopy images of 786-O cells transfected with empty vector (CT-GFP) or GFP-tagged *PCDHGC3* vector (GFP-GC3) 48-hours post-transfection. White arrows point to GFP staining localized at the cell membranes and cell-to-cell contacts. (E) Percentage of GFP-expressing cells determined by flow cytometry analysis 24 hours post-transfection. (F) Cell viability assessed by MTS assay in 786-O cells transfected with CT-GFP and GFP-GC3 at different time points after transfection. (G) Flow cytometry-based cell cycle analysis of the indicated cells at 12-, 24- and 36-hours post-transfection. (H) Schematic representation of the mouse xenograft model and representative tumor images from CT and C3KD xenografts. (I) *PCDHGC3* mRNA levels in CT and C3KD tumor xenografts (n = 3 tumors per condition) (J) Volume (left) and weight (right) of tumors derived from CT or C3KD 786-O cells. Data represent mean value ± SEM. n = 9 tumors per group. (K) Representative images of hematoxylin & eosin (H&E) staining and Ki67 immunohistochemistry of CT and C3KD tumor xenografts. The bar plot represents the percentage of Ki67 positively stained cells. *p < 0.05, **p < 0.01, *** p < 0.001, **** p< 0.0001; Scale bars = 200 μm.

To assess the persistence of these effects under more physiological conditions, we examined cell growth in a three-dimensional (3D) environment using our newly developed 3D bioprinted cancer model (Herrada-Manchón *et al*, 2021). This model involves printing cells in a mixture of collagen, sodium alginate, and gelatin. As reported previously, the printed cells remained viable and able to grow for up to 15 days. Over a 14-day period, we found a significant increase in cell growth in C3KD cells compared to CT cells, with a 1.33- or 2.82-fold increase in 786-O and RCC4 cells, respectively (Fig 2C).

We then sought to examine the potential influence of *PCDHGC3* over-expression on cell proliferation. Initially, we transiently transfected 786-O cells with GFP-tagged *PCDHGC3* to induce overexpression of this gene. GFP expressing vector was used as a control (CT-GFP). Subsequent microscopy analysis verified the effective recruitment and clustering of the exogenous fusion protein at both cell membranes and cell-to-cell contact sites (Fig 2D). As shown in Fig 2E, the transfection efficiency was relatively low (approximately 15-20% of cells) as determined by flow cytometry analysis of GFP-positive cells, revealing no significant differences between CT-GFP and GFP-*PCDHGC3* transfected cells. To assess cell growth, we employed MTS assays, which displayed no significant alteration in GFP-tagged *PCDHGC3* compared to CT-GFP cells (Fig 2F), likely due to the modest transfection efficiency. Our previous findings demonstrated a 20% increase in cells within the S phase of the cell cycle upon reduced *PCDHGC3* expression (Bernardo-Castiñeira *et al*, 2019). Consequently, we analyzed the cell cycle distribution of cells overexpressing GFP-*PCDHGC3*. Despite the modest transfection efficiency, our analysis of the cell cycle in transduced GFP-expressing cells unveiled a noteworthy 15% reduction in cells within the S phase following *PCDHGC3* overexpression (Fig 2E, G), thereby providing further evidence of its role in modulating cell proliferation.

We then investigated whether *PCDHGC3* knockdown increases ccRCC tumor growth *in vivo*. We used a xenograft-based mouse model of ccRCC by subcutaneous injection of C3KD and CT cells into immunocompromised mice (Fig 2H). Surprisingly, neither C3KD- nor CT-RCC4 cells formed tumors during the 60-day observation period, unlike 786-O cells, both C3KD and CT, that did form tumors (Fig 2H). The maintenance of *PCDHGC3* silencing was confirmed in these tumors (Fig 2I). Notably, we observed that tumors originating from C3KD-cells exhibited significantly higher volume (4.1-fold increase) and weight (5.5-fold increase) compared to those from CT-cells over the 60 days period (Fig 2J). Additionally, C3KD-derived tumors displayed a higher percentage of Ki67-positive cells (51 ± 16.73%) compared to CT cells (30.16 ± 8.54%) indicative of increased cellular proliferation (Fig 2K).

### *PCDHGC3* knockdown induces EMT, cell survival and metastatic growth

To explore the potential role of *PCDHGC3* in metastasis, we first revisited the xenograft-mice previously generated. No metastases were identified in kidney, lung, liver, and brain in any of the mice. However, evidence of cancer cell infiltration in the dermis was detected in two animals xenografted with C3KD but not CT cells (Fig 3A). Thus, we explored the putative involvement of *PCDHGC3* in the epithelial-mesenchymal transition (EMT) pathway which plays a crucial role in local cell invasion. As depicted in Fig 3B and C, western blot analysis revealed reduced levels of the epithelial marker cytokeratin and increased expression of mesenchymal markers such as N-cadherin, ZEB1, ZEB2, and Snail2 in C3KD cells compared with CT cells. These changes suggest that *PCDHGC3* knockdown promotes the progression of EMT.

**Figure 3.**
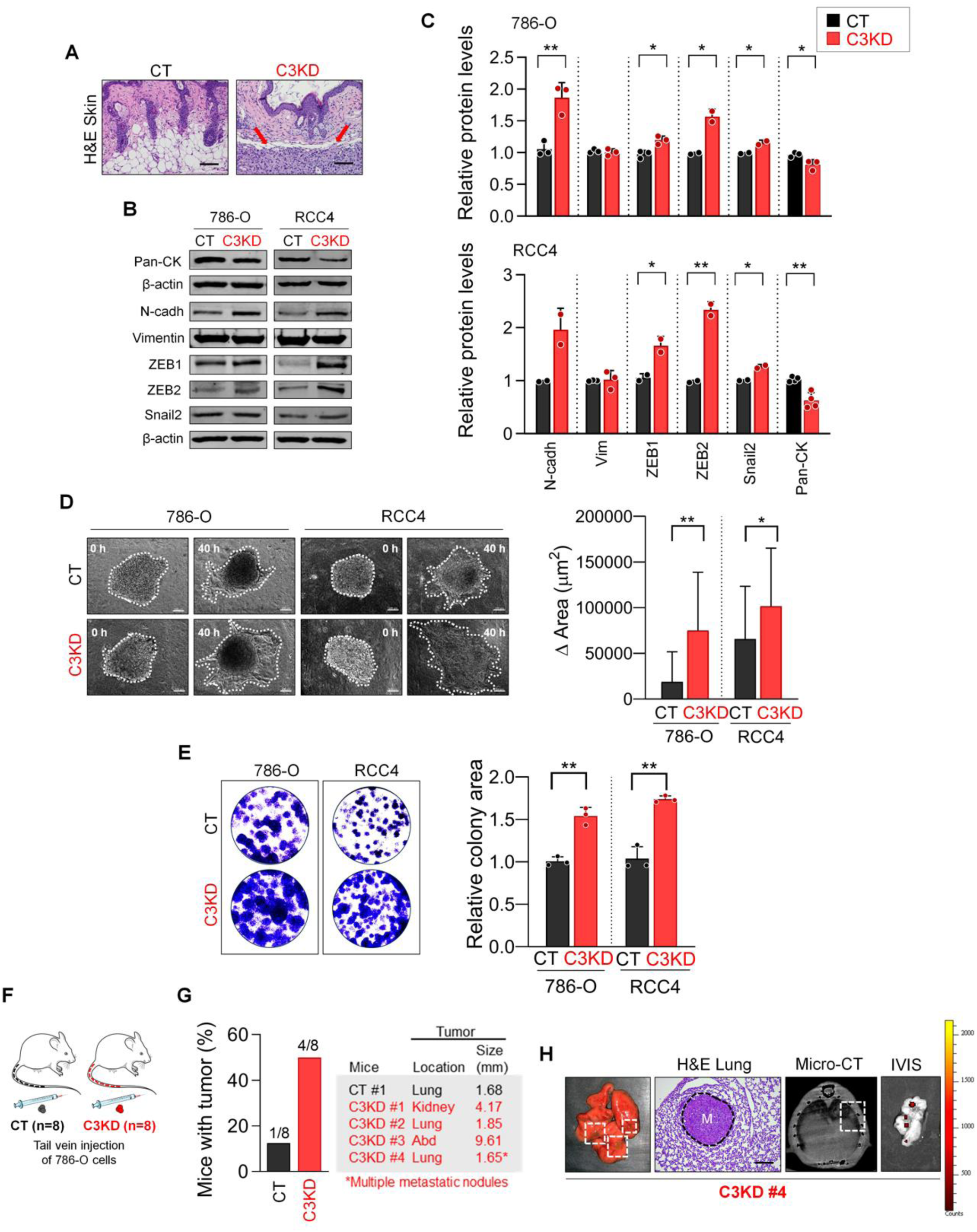
*PCDHGC3* knockdown promotes metastatic cell phenotypes. (A) Representative images of hematoxylin & eosin (H&E) staining of skin adjacent to CT and SH 786-O tumor xenografts. (B, C) Immunoblot of pan-cytokeratin (pan-CK) or the indicated mesenchymal biomarkers in CT and C3KD cells. Expression of anti-β-actin was used for normalization. Bar plots show the densitometric quantification of the relative protein levels. (D) Images show 786-O or RCC4 CT/C3KD cell spheroids assembled after seeding cells on top of bioink surfaces (0 h) and incubated for 40 h. The graph represents variations in spheroid invasion area over a 40-hour period. For area quantification, at least 8 spheroids from a minimum of 2 independent experiments were used. Data are represented as mean ± SD. Scale bars: 100μm. (E) Representative images and quantification of relative colony areas formed by 786-O and RCC4 CT and C3KD cells measured by cell colony formation assay. (F) Schematic representation of the experimental design. (G) Frequency, location, and size of tumors developed from CT and C3KD cells injected into the tail vein of mice and detected by Micro-CT scans 4 months post-inoculation. (H) Representative images of H&E staining, Micro-CT scan, and In Vivo Imaging System (IVIS) of green fluorescence in a C3KD-derived lung tumor. *p < 0.05, **p < 0.01.

Subsequently, we examined the effect of silencing *PCDHGC3* on the different stages of the metastatic cascade: local invasion, survival in hostile environments, and growth in distant organs.

Cell invasion was analysed in vitro by seeding cells onto pre-printed matrices. Consistent with previous findings (Herrada-Manchón *et al*, 2021), the cells exhibited spontaneous formation of multicellular spheroids after a 24-hour incubation period, subsequently adhering to and invading the matrices. Using our in-house software, we tracked spheroid areas over time to assess cell invasive potential. Our analysis revealed a significant increase in invasive potential of C3KD 786-O (3.98-fold, p = 0.0023) and RCC4 (1.6-fold, p = 0.03) cells compared to CT cells (Fig 3D).

We previously showed that the knockdown of *PCDHGC3* led to an increased clonogenic capacity of 786-O cells (Bernardo-Castiñeira *et al*, 2019). Expanding on this, we explored cell survival in challenging environments using colony formation assays in both CT and C3KD-RCC4 cells. Our findings revealed that C3KD cells exhibited a superior survival rate and sustained growth in comparison to CT cells (1.53- and 1.68-fold increase in 786-O and RCC4 cells, respectively) (Fig 3E). To directly assess the capacity of *PCDHGC3*-deficient cells to survive in the bloodstream and spread to diverse organs, we thoroughly examined the whole bodies of mice 4 months after the intravenous injection of C3KD or CT 786-O cells (Fig 3F). Micro-computed tomography (CT) scans revealed that 4 out of 8 mice injected with C3KD cells developed tumors in different organs, while only 1 out of 8 mice injected with CT cells showed tumor growth (Fig 3G, H). Subsequent histological analysis of these tissues, along with IVIS Spectrum In Vivo Imaging to detect TurboGFP-labeled cells, confirmed the presence of tumor cells in the target organs (Fig 3H).

### Activation of the mTOR and HIF signaling pathways in *PCDHGC3* deficient cells

To identify the mechanisms by which *PCDHGC3* inhibition increases tumor growth and invasion, we examined HIF, PI3K-mTOR, and ERK pathways, recognized as pivotal in the pathogenesis of ccRCC and as potential therapeutic targets (Ahmed & Ornstein, 2023),(González-Larriba *et al*, 2017),(Liu *et al*, 2016).

Through Western blot analysis, we found increased levels of phosphorylated mTOR (Ser2448 p-mTOR), AKT (Ser473 p-AKT), ribosomal protein S6 (Ser240/244 p-S6), and 4EBP1 (Thr37/46 p-4E-BP1), which are regulators activated by the PI3K-mTOR pathway (Fig 4A, B). Treating with specific mTOR inhibitors, rapamycin and temsirolimus, effectively reduced these levels (Fig 4C, D). Additionally, C3KD cells showed elevated levels of phosphorylated Erk1/2 (Thr202/204 p-Erk1/2), potentially regulating the mTOR pathway, compared to CT cells. However, there were no significant changes in phosphorylated GSK-3β (Ser9 pGSK-3β), a component of the Wnt signaling pathway. Overall, these findings strongly suggest that *PCDHGC3* has a significant impact on the mTOR and Erk pathways. Nonetheless, the role of *PCDHGC3* in the Wnt signaling pathway remains to be further explored.

**Figure 4.**
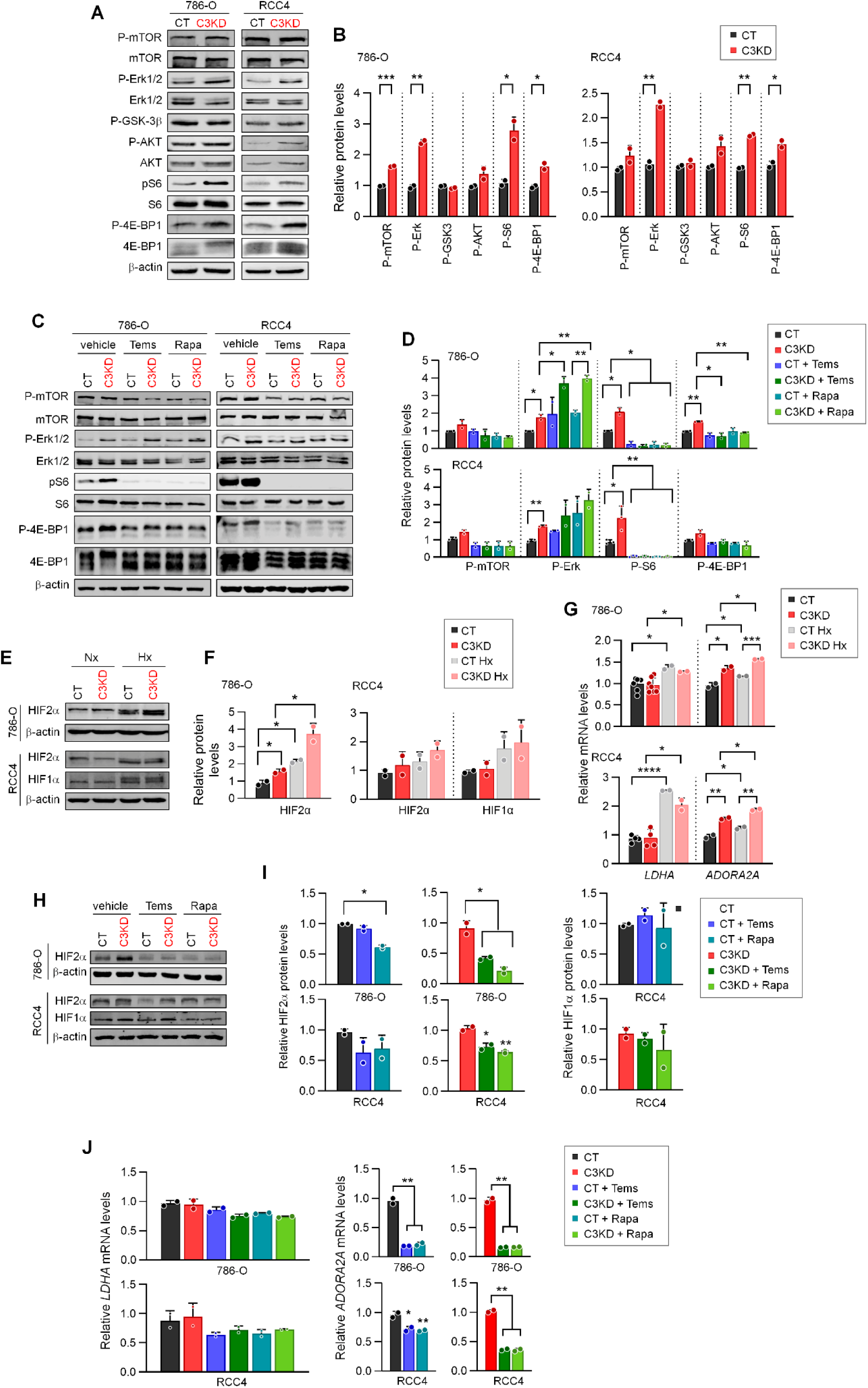
*PCDHGC3* knockdown induces activation of mTOR-HIF2α signaling. (A-F, H-I) Representative immunoblots images and quantifications of the indicated proteins. Cells were treated with 1 μM temsirolimus (Tems) or rapamycin (Rapa) for 24 hours. Hypoxia was induced by incubating cells at 1% O2 for 12 hours. All immunoblot data are normalized to β-actin levels. (G, J) Relative mRNA levels of the indicated HIF target genes in 786-O and RCC4 CT and C3KD cells. Nx: normoxia, Hx: hypoxia. *p < 0.05, **p < 0.01, *** p < 0.001. **** p < 0.0001.

Next, we analyzed the protein levels of HIF1α and HIF2α, key transcription factors in ccRCC development (Kaelin, 2007) (Fig 4E, F). In 786-O cells, only HIF2α is expressed, while RCC4 cells express both, HIF1α and HIF2α (Iliopoulos *et al*, 1995). The two cell lines accumulate these factors even under normal oxygen levels due to dysfunctional pVHL (Fig 4E, F). Our results confirmed that both HIF1α and HIF2α increased under low oxygen conditions (1% O2, 12 hours), as expected. Additionally, HIF2α levels were higher in C3KD-786-O cells than in CT cells even under normal oxygen conditions, with a less noticeable effect in RCC4 cells (Fig 4E, F). Interestingly, reducing *PCDHGC3* levels did not consistently change HIF1α levels in RCC4 cells (Fig 4E, F). Moreover, *LDHA*, a gene regulated mainly by HIF1α, did not show increased expression in C3KD cells (Fig 4G). However, the expression of *ADORA2A*, a gene targeted by HIF2α, significantly increased in both C3KD-786-O and C3KD-RCC4 cells as compared to CT cells (Fig 4G).

Our data also revealed that the mTOR inhibitors counteracted the impact of *PCDHGC3* knockdown on HIF2α, while not affecting HIF1α levels (Fig 4H, I). This observation suggests an mTOR-mediated mechanism specifically regulating HIF2α. Furthermore, the expression of *ADORA2A*, but not *LDHA*, was also found to be mTOR-dependent as revealed by its reduced expression following exposure to temsirolimus and rapamycin (Fig 4J). Notably, these regulatory effects were markedly more pronounced in C3KD cells compared to CT cells.

We also confirmed the sustained effect of *PCDHGC3* knockdown on mTOR and HIF2α pathways in the tumor xenografts by the analysis of the signaling proteins in tumor tissue samples (Fig EV2). Additionally, immunohistochemical analysis revealed higher expression of HIF2α in C3KD compared to CT xenografts, with precise localization within tumor cell nuclei. However, contrary to findings in cell lines, the increased activation of Erk1/2 was not observed in C3KD xenografts.

### The role of mTOR and HIF pathways in regulating cell growth and survival in *PCDHGC3*-deficient cells

To understand if the increased cell proliferation due to *PCDHGC3* knockdown relies on mTOR or HIF2α, we treated cells with maximum inhibitory concentrations of rapamycin, temsirolimus, or the HIF2α inhibitor PT2385.

Cell growth of 786-O cells was measured three days after treatment using either the MTS assay (Fig 5A) or continuous monitoring over a 90-hour period with the iCelligence system (Fig 5B). Results from both methods indicated that both temsirolimus and rapamycin effectively inhibited cell proliferation in both C3KD and CT 786-O cells, with a more pronounced effect observed in C3KD cells, whose proliferation was reduced to a greater extent (1.22 to 1.38-fold) than that of CT cells (p<0.04). Similar results were observed in RCC4 cells (Fig EV3). Additionally, PT2385 exhibited no effect on CT cells proliferation but demonstrated a significant reduction of 1.32-fold (p<0.0001) in the growth of C3KD 786-O cells. Combining mTOR and HIF2α inhibitors revealed no further difference from each alone, suggesting a shared pathway inhibited by these compounds (Fig 5A). Cell growth was further evaluated within the 3D bioprinted cancer model, revealing that the impact of PT2885 and mTOR inhibitors in the 3D setup closely paralleled the observations made in traditional 2D culture (Fig 5C, D).

**Figure 5.**
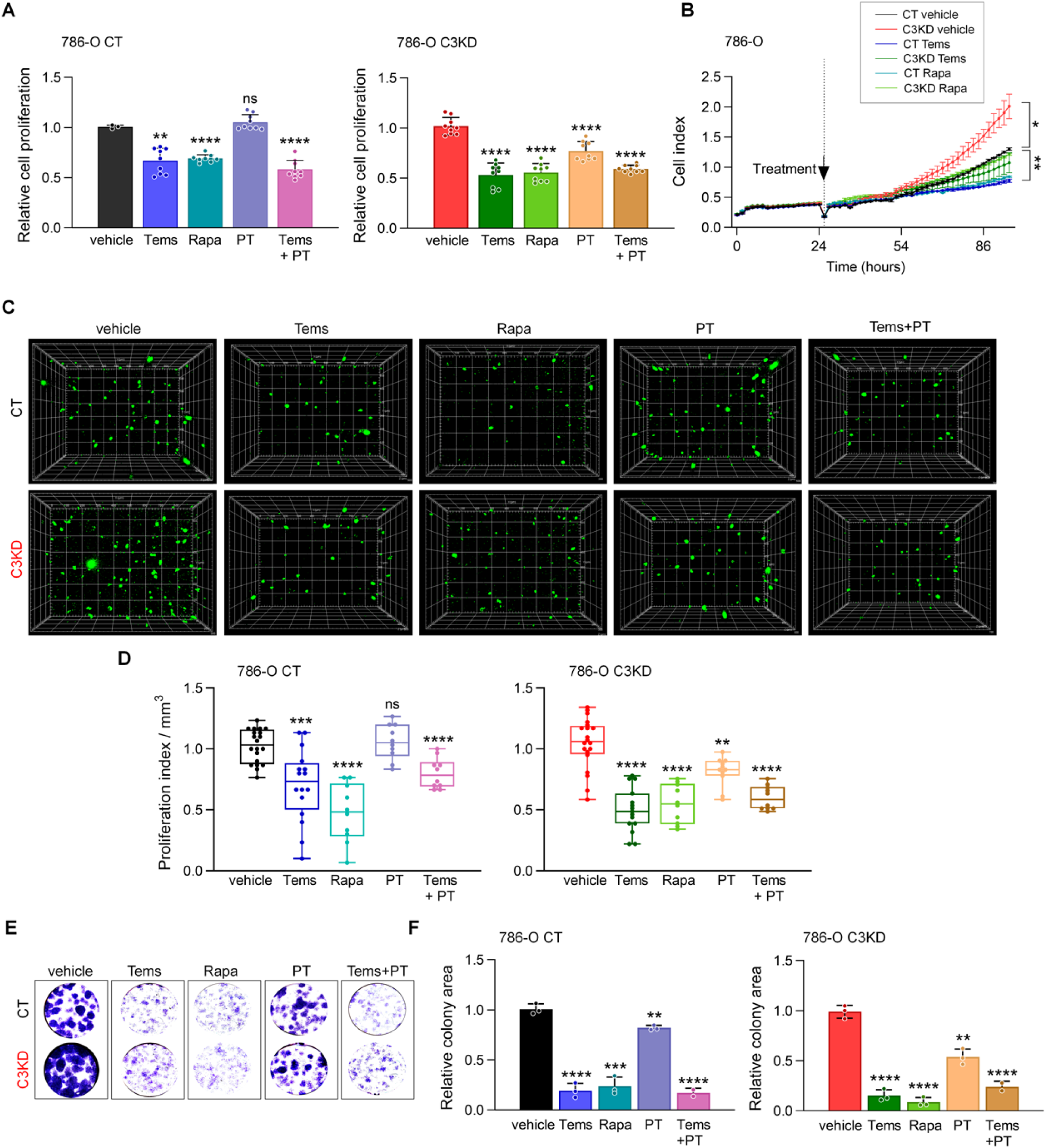
mTOR and HIF2α inhibitors suppress high proliferative rate induced by *PCDHGC3* deficiency in ccRCC cells. (A) MTS cell proliferation of the indicated cells 72 hours after treatment with 1 μM temsirolimus (Tems), 1 μM rapamycin (Rapa) or 10 μM PT2385 (PT). (B) Real-time analysis of cell proliferation using the iCELLigence system in 786-O CT or C3KD cells treated or not with temsirolimus or rapamycin (1 μM). (C) Representative fluorescence XYZ projections of 786-O CT or C3KD printed cells 786-O treated with the indicated drugs. (D) Quantification of bioprinted cells after treatment with different drugs. Cell counting was performed on Z-stack images (15-20 sections with an interval of 20 μm between them) at randomly chosen positions within the scaffold. (E, F) Representative images and quantification of relative colony areas formed by CT and C3KD 786-O cells measured by colony formation assay. DMSO was used as vehicle in all assays. Data show mean value ± SD (n ≥ 3 independent experiments). *p < 0.05, **p < 0.01, *** p < 0.001, **** p < 0.0001.

Our subsequent colony assays also revealed that mTOR inhibitors had a more pronounced effect in C3KD compared to CT cells (1.7-2.9-fold greater reduction, p<0.05) (Fig 5E, F). Likewise, PT2385 either exhibited no effect or had a minimal impact on CT cells, while significantly reducing colony formation in C3KD cells. Similar results were obtained in RCC4 cells (Fig EV3).

### Therapeutic potential of temsirolimus in *PCDHGC3*-deficient tumors

Overall, the data indicate that *PCDHGC3* knockdown enhances tumor growth and cell invasion, survival, and spread. Importantly, our results demonstrate that these effects can be reversed by various drugs, including temsirolimus. To evaluate the effectiveness of temsirolimus as a treatment for tumors with and without normal *PCDHGC3* expression, we employed two distinct mouse models.

In a xenograft model, we started temsirolimus treatment 66 days after subcutaneous inoculation of either CT or C3KD 786-O cells, to align with the time required for tumor formation (Fig 6A). The treatment regimen consisted of two cycles, each lasting 5 days, with a 2-day rest interval between cycles. Tumor volume and weight assessments post-treatment indicated a partial response to temsirolimus (Fig 6B-D). A comparable effect was observed in C3KD xenografts (with a range of 1 to 26-fold reduction in tumor weight) and CT xenografts (with a range of 5 to 11-fold reduction in tumor weight) (p=0.990). Throughout the treatment process, *PCDHGC3* levels remained unchanged, as confirmed by analyzing *PCDHGC3* expression in both control and treated tumors (Fig 6E). Histological examination of tumor tissue revealed significant fibrosis in treated tumors from both CT and C3KD xenografts, with the additional observation of increased necrosis in C3KD xenografts (Fig 6F). Ki67 levels indicated higher proliferation activity in C3KD xenografts, with minimal changes evident after treatment in both CT and C3KD xenografts (Fig 6F). Therefore, despite the observed increase in necrosis in C3KD xenografts, according to this mouse model, *PCDHGC3* deficient tumors did not show enhanced vulnerability to temsirolimus treatment.

**Figure 6.**
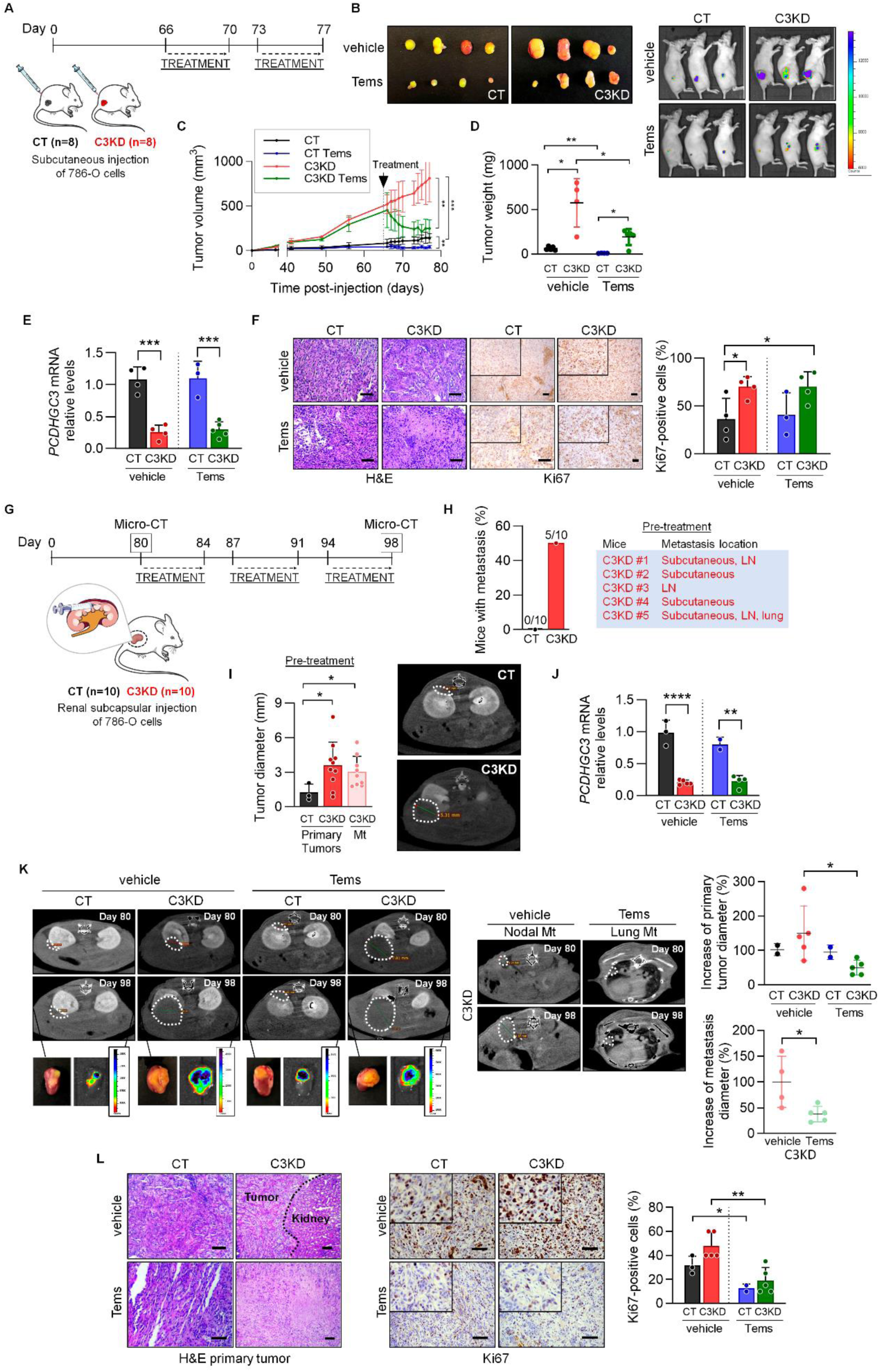
mTOR inhibition reduces increased tumor growth and blocks metastasis development triggered by *PCDHGC3* silencing in vivo. (A-F) Tumor xenograft model. (A) Schematic overview of the experimental procedure in the mouse xenograft model. Vehicle (DMSO) or temsirolimus (Tems) were administered intraperitoneally 66 days after cell inoculation in athymic nude mice at a dosage of 1.5 mg/kg/day, 5days/week for 2 weeks. (B) Representative images of tumor xenografts and IVIS detection of green fluorescence in the whole animals after treatments. Tumor volume (C) and weight (D) of the indicated tumor xenografts under the specified treatments. Data represent mean value ± SEM. (E) Relative *PCDHGC3* mRNA levels measured in xenograft tumors after treatments. (F) H&E staining and Ki67 immunohistochemical images of the indicated tumors xenografts. (G-L) Orthotopic model. (G) Flowchart of the experimental design. Vehicle (DMSO) or temsirolimus (1.5 mg/kg/day) treatments were administered intraperitoneally 5 days a week for 3 weeks. (H) Pretreatment analysis by Micro-CT showing the percentage of mice with metastasis and the metastasis location. LN: lymph nodes. (I) Bar chart represents the mean diameter (mm) ± SD of primary tumor and metastases (Mt) measured 80 days after cell inoculation by analysis of Micro-CT. Representative images of Micro-CT scans of the indicated tumors before treatments are shown. (J) *PCDHGC3* mRNA levels in the indicated tumors after treatments. (K) Representative Micro-CT scans of mice before (Day 80) and after (Day 98) treatment showing primary tumors and metastases. Response to treatments is shown as the variation of tumor diameter between day 80 and 98 in mice treated with vehicle or temsirolimus. (L) Ki-67 immunohistochemical and H&E stainings of the indicated primary tumors. Scale bars = 200 μm. *p < 0.05, **p < 0.01, *** p < 0.001. **** p < 0.0001.

While the xenograft model proved valuable in assessing the effects of *PCDHGC3* and temsirolimus on tumor growth, its capability to study metastasis development was limited, as subcutaneous tumors did not metastasize. Acknowledging that orthotopic xenografts better replicate the native tumor microenvironment and more accurately mimic the growth and metastatic behavior of human tumors compared to subcutaneous xenografts (Zheng, 2019; Lwin *et al*, 2018), we subsequently transitioned to an orthotopic mouse model. In this model, cells were injected into the subcapsular region of the kidney, followed by the treatment regimen outlined in Fig 6G. Before treatment, whole-body Micro-CT scans showed tumors in all animals. However, while none of the control group’s animals had metastasis, 5 out of 10 mice injected with C3KD cells developed lymph node, subcutaneous, or lung metastases by day 80 (Fig 6H). In addition, C3KD primary tumors were notably larger than CT tumors (2.93-fold, p=0.01) (Fig 6I).

Temsirolimus treatment began on day 80, with three 5-day cycles and 2-day rest intervals in between. The sustained suppression of *PCDHGC3* was verified in the primary tumors at the end of the experiments (Fig 6J). Remarkably, both primary C3KD tumors and their metastases showed a significant response to temsirolimus, with 67% and 62% reductions in growth, respectively (Fig 6K). Interestingly, no response to temsirolimus was noted in CT tumors. Histological examination confirmed earlier findings of high fibrosis and necrosis in temsirolimus-treated tumors (Fig 6L). We also found higher Ki67 levels in C3KD tumors than in CT tumors, with a notable decrease after temsirolimus treatment (Fig 6L). Furthermore, temsirolimus reduced p-S6, p-4E-BP1, and HIF2α levels significantly in both CT and C3KD primary tumors, as well as in C3KD-derived metastases (Fig EV4).

Collectively, the data indicate that the knockdown of *PCDHGC3* rendered cancer cells highly susceptible to mTOR inhibition, exhibiting marked sensitivity in terms of both tumor growth and metastasis development.

### Lipid metabolism reprograming induced by *PCDHGC3* silencing

To comprehensively decipher the signaling pathways activated in C3KD cells, we performed a quantitative proteomic analysis. We compared 5 replicates each of C3KD and CT 786-O cells, identifying 7350 protein groups representing 7230 quantifiable proteins. Among them, 292 (4%) showed significant upregulation, while 458 (6%) were significantly downregulated in C3KD cells compared to CT cells (fold change = 1.2, Benjamini-Hochberg adjusted *p* < 0.01, limma test) (Fig 7A and Table EV1).

**Figure 7.**
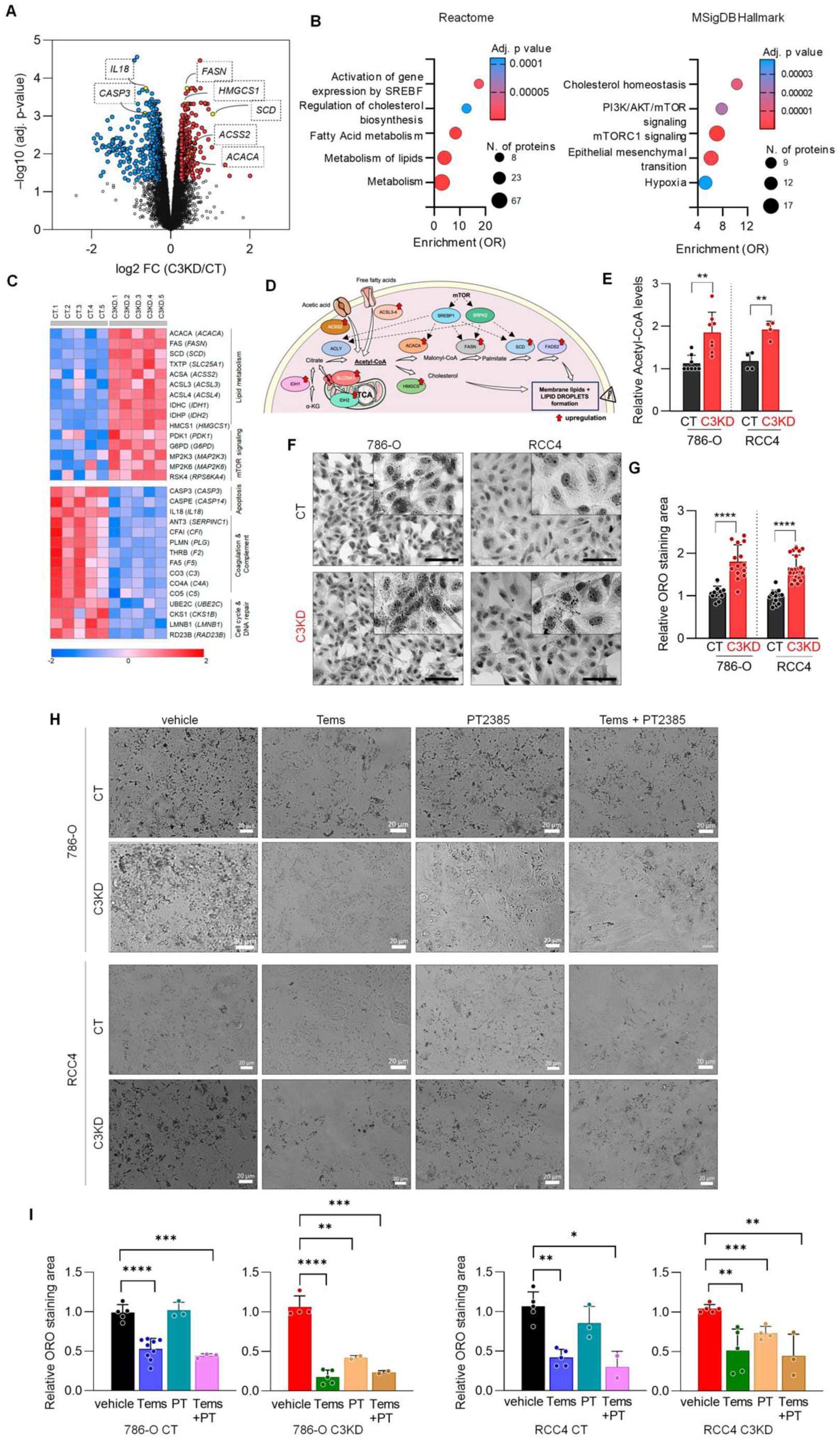
Lipid metabolism reprograming following *PCDHGC3* knockdown. (A) Volcano plot representation of proteins identified by proteomic analysis of C3KD vs. CT 786-O cells. Proteins significantly up- or down-expressed (adjusted p value < 0.01 and > 1.2-fold-cutoff) are highlighted by red or blue circles, respectively. (B) Enrichment analysis of up-regulated proteins using Reactome and Molecular Signature (MSigDB) Hallmarks databases. (C) Heatmap visualization of the most significantly differentiated proteins. (D) Schematic representation of the pathways involved in lipid and cholesterol synthesis. Red arrows indicate proteins upregulated in 786-O C3KD cells compared to CT cells. (E) Relative intracellular acetyl-CoA levels in CT and C3KD cells. (F-H) Representative images of phase-contrast microscopy of the indicated cells stained with Oil Red O (ORO) three days after reaching confluence. Temsirolimus (1 μM) and PT2385 (50 μM) treatments were performed for 72 hours once cells reached confluence. (G, I) Bar charts show quantification of ORO area staining. Scale bars = 100 μm (F) and 20 μm (H). *p < 0.05, **p < 0.01, **** p < 0.0001.

Functional enrichment analysis of differentially expressed proteins confirmed higher levels of proteins associated with mTORC1, HIF, and EMT pathways in C3KD cells than in CT cells (Fig 7B and Table EV2). On the other hand, pathways like complement and coagulation cascades, apoptosis, cell cycle regulation, and DNA repair showed significant downregulation in C3KD cells (Fig 7C and Table EV2). Metabolic pathways showed a notable increase among upregulated proteins in C3KD cells. However, we did not find any impact of *PCDHGC3* silencing on mitochondrial function when assessed with Cell Mito Stress Seahorse assay (Fig EV5). The enrichment of lipid metabolism, especially fatty acid (FA) biosynthesis, was noteworthy, involving key proteins such as SLC25A1, ACSS2, ACSL3, ACACA, FASN, SDC, and FADS2 (Fig 7D and Table EV2). This suggests their potential role in abnormal pathway activation linked to acetyl-CoA and FA synthesis. Acetyl-CoA levels were found significantly higher in C3KD cells compared to CT cells (1.7-fold, p < 0.002, Fig 7E), indicating a potential imbalance leading to increased storage of FA in cytoplasmic lipid droplets. Staining with Oil Red O confirmed a notable increase in lipid droplets in C3KD cells (1.7-fold, p < 0.0001, Fig 7F and 7G).

Treatment with temsirolimus substantially reduced lipid droplets, particularly in C3KD 786-O cells (Fig 7H, 7I). Interestingly, inhibition of HIF2α with PT2385 had no effect on CT cells but significantly reduced lipid droplet density in both 786-O and RCC4 C3KD cells. This effect was more pronounced in 786-O cells, which accumulate HIF2α but not HIF1α, compared to RCC4 cells that express both HIFα subunits. Treatment with both temsirolimus and PT2385 did not demonstrate a greater effect than either compound alone. These findings suggest that lipid droplet accumulation in C3KD cells is primarily mediated by HIF2α activity.

## Discussion

This study unveils a significant advancement in the understanding of the pathogenesis of metastatic ccRCC. We reveal, for the first time, the impact of *PCDHGC3* downregulation on tumor growth, cancer cell local invasion, and metastasis development.

PCDHs, belonging to the cadherin superfamily, are cell adhesion molecules crucial for intercellular adhesion and cell-cell communication, key mechanisms of tumor growth and metastasis. Although the epigenetic silencing of *PCDHG* genes has been identified in various types of cancer and appears to be involved in tumorigenesis, particularly in colon cancer (Dallosso *et al*, 2012b), the role of an individual *PCDHG* gene in metastasis has not been previously documented, to the best of our knowledge. In our previous studies, we reported an association between *PCDHGC3* methylation and the aggressive and metastatic forms of paraganglioma, pheochromocytoma (Bernardo-Castiñeira *et al*, 2019) and gastrointestinal neuroendocrine carcinoma (Cubiella *et al*, 2024). Here, our in vitro and in vivo investigations reveal the substantial involvement of the *PCDHGC3* gene in every stage of metastasis suggesting its potential role as a tumor suppressor gene. Notably, our in-silico analysis establishes a compelling correlation between *PCDHGC3* silencing and advanced disease stage and reduced overall disease survival in ccRCC patients. Collectively, these data underscore the significance of *PCDHGC3* in the pathogenesis of ccRCC.

While several roles of *cPCDHs* in neuronal development have been mechanistically dissected (Peek *et al*, 2017), the molecular processes they mediate in cancer have remained mostly unknown. In general, *cPCDH* proteins interact with a range of molecules to regulate diverse downstream signaling pathways, including Wnt/β-catenin, mTOR and FAK tyrosine kinases (Chen *et al*, 2009; Keeler *et al*, 2015b; Dallosso *et al*, 2009). Crucially, our research identifies the mTOR and HIF2α pathways as downstream signaling targets influenced by *PCDHGC3* in ccRCC cancer cells. It is noteworthy that both the mTOR and HIF2α signaling pathways are hyperactivated in most ccRCC. The mTOR pathway functions as a metabolic, cell proliferation, and survival regulator, and its activity, which is essential for tumor growth and progression, can be pharmacologically targeted. Particularly, mTOR inhibitors, such as temsirolimus, stand as the established standard treatment for patients with metastatic ccRCC (González-Larriba *et al*, 2017). Additionally, the recently developed HIF2α inhibitor, Belzutifan (the second-generation small-molecule HIF2α inhibitors), emerges as a highly promising therapeutic option for metastatic ccRCC, demonstrating noteworthy efficacy, particularly in cases associated with VHL disease(Ahmed & Ornstein, 2023; Nguyen *et al*, 2024; Jonasch *et al*, 2024; Choueiri *et al*, 2023; Chan *et al*, 2024). In our study, we illustrate that both mTOR and HIF2α inhibitors counteract the positive effects of *PCDHGC3* silencing on tumor growth and metastasis. Moreover, the combination of these two drugs does not exhibit a cumulative inhibitory effect, indicating their impact on the same signaling pathway. Consistently, we found that the *PCDHGC3*-dependent activation of HIF2α is dependent on mTOR activity. Dependence of HIF2α on mTORC1 in ccRCC cells has been previously demonstrated (Toschi *et al*, 2008). Collectively, these findings point towards the potential for personalized medicine, where patients with *PCDHGC3* deficiencies may benefit more effectively from the use of mTOR or HIF2α inhibitory drugs.

Proteomic and metabolomic analyses further reveal that *PCDHGC3* silencing has no discernible impact on mitochondrial activity but induces the overexpression of proteins related to FA synthesis. These include acetyl-CoA carboxylase (ACACA) (a rate-limiting enzyme in FA synthesis), FA synthase (FASN) (a major lipogenic protein), and the desaturases SCD or FADS2, integral to FA synthesis and direct targets of the sterol regulatory element-binding transcription factor 1 (SREBP1) activated by mTOR, along with hidroximetilglutaril-CoA synthase (HMGCS) involved in cholesterol synthesis. Additionally, *PCDHGC3* downregulation correlates with elevated levels of ATP citrate lyase (ACLY) and acyl-CoA synthetase (ACSS2) proteins and transcripts, contributing to acetylCoA synthesis, a precursor metabolite in FA biosynthesis. Notably, acetylCoA significantly accumulates in *PCDHGC3*-deficient cells compared to *PCDHGC3*-proficient cells. Moreover, our findings indicate that *PCDHGC3* silencing leads to increased lipid droplets, a process modulated by the mTOR pathway as suggested in previous reports (Zhang *et al*, 2024; Mensah *et al*, 2017; Fuentes *et al*, 2019). Nevertheless, we observed that the HIF2α inhibitor, PT2385, does not affect lipid droplet density in CT cells but significantly reduce it in C3KD cells. Previous studies have shown that increased de novo FA synthesis and uptake, which eventually leads to lipid droplet formation, are positively regulated by *VHL* loss of function and subsequent HIF2α activation in ccRCC (Qiu *et al*, 2015; Zhang *et al*, 2024). Thus, the decreased expression of *PCDHGC3* not only activates both mTOR and HIF2α but also significantly amplifies FA synthesis, a process that can be mostly attributed to the HIF2α signaling.

The regulation of essential metabolic pathways underscores a potent mechanism through which *PCDHGC3* deficiency actively promotes tumor metastasis and overgrowth. Reprogramming of lipid metabolism is a characteristic feature of ccRCC, correlating with tumor aggressiveness and poor prognosis. Specifically, many FA enzyme alterations are known to occur in ccRCC in correlation with poor clinical outcomes (Tan *et al*, 2023). Particularly noteworthy is the role of lipid metabolism in the metastatic cascade (Martin-Perez *et al*, 2022). Cells with metastatic capacity exhibit increased lipid metabolism and expression of FA transporters (Ladanyi *et al*, 2018; Pan *et al*, 2019; Jiang *et al*, 2019). Moreover, enhanced FA uptake favors the invasion and migration processes triggered by the EMT of metastatic-initiating cells (Corbet *et al*, 2020). Finally, increased de novo lipid biosynthesis through ACLY and FASN, promotes the migration and invasion abilities of cancer cells (Wen *et al*, 2019; Guo *et al*, 2019).

Our results also contribute to the mounting evidence that individual clustered protocadherin family members can have unique biological functions both inside and outside of the nervous system. The identification of the cPCDH loci (Wu & Maniatis, 1999), the elucidation of mechanisms controlling the semi-stochastic expression of their >50 individual gene isoforms (Tasic *et al*, 2002; Wang *et al*, 2002; Ribich *et al*, 2006; Kaneko *et al*, 2006; Kawaguchi *et al*, 2008; Guo *et al*, 2012; Canzio *et al*, 2019), and the demonstration that clustered protocadherin proteins promiscuously form *cis-*dimers that interact strictly homophilically in *trans* (Schreiner & Weiner, 2010; Thu *et al*, 2014; Rubinstein *et al*, 2015; Goodman *et al*, 2017) all pointed to the importance of isoform diversity in clustered protocadherin function. Recently, however, complex genetic analyses in mice have demonstrated distinct roles for individual broadly expressed C-type proteins in neural development that cannot be duplicated by other isoforms. For example:

α-Pcdh-C2 is uniquely critical for the projection of serotonin-producing neurons within multiple brain regions (Chen *et al*, 2017); γ-Pcdh-C3 is particularly important for dendritic arborization and synapse formation (Steffen *et al*, 2023; Meltzer *et al*, 2023); and γ-Pcdh-C4 is the only clustered Pcdh protein strictly required for neuronal survival (Garrett *et al*, 2019; Leon *et al*, 2024). The emergence of *PCDHGC3* as a possible tumor suppressor gene in a wide variety of cancers, now including ccRCC, bolsters the emerging conclusion that some biological clustered protocadherin functions are controlled by a single isoform, and thus do not require family diversity.

How might individual clustered protocadherin proteins mediate such unique functions? One likely possibility is that they engage unique combinations of intracellular signaling proteins via their unique membrane-proximal variable cytoplasmic domains (VCDs). Such signaling partners remain unknown for most clustered protocadherin proteins, but recent work has identified Axin1, a scaffolding protein and Wnt signaling component, as a unique binding partner of the γ-Pcdh-C3 VCD (Steffen *et al*, 2023; Mah *et al*, 2016). This interaction mediates the isoform’s unique ability to inhibit canonical Wnt signaling *in vitro* (Mah *et al*, 2016). Interestingly, Axin1 itself has been reported to act as a tumor suppressor and can regulate mTOR signaling (Qiu *et al*, 2024). Future structure-function analyses of the human PCDHGC3 protein may determine the extent to which signaling through its VCD is required for its role in ccRCC, either through Axin1 or other unknown binding partners.

In conclusion, our study unveils, for the first time, the significant impact of a clustered protocadherin gene on ccRCC growth and metastasis through the activation of the mTOR-HIF2α pathway and the reprogramming of lipid metabolism. This highlights a potential mechanism underlying the aggressive behavior of ccRCC, underscoring the promise of personalized therapeutic approaches. Specifically, patients with *PCDHGC3* deficiency may benefit more effectively from mTOR- or HIF2α-inhibitory drugs, as well as from small-molecule inhibitors targeting the FA biosynthetic pathway.

## Methods

### Cell cultures

Clear cell renal cell carcinoma 786-O and RCC4 cell lines were kindly provided by Dr. M.J. Calzada (Hospital Universitario de La Princesa, Madrid, Spain). Cells were cultured in Dulbecco’s Modified Eagle Medium (DMEM) supplemented with 10% v/v heat-inactivated Fetal Bovine Serum (Gibco, Thermo Fisher Scientific), 100 U/mL penicillin, and 100 μg/mL streptomycin, and maintained at 37⁰C, 5% CO2. During hypoxia experiments, cells at 70-80% confluence were exposed to 1% O2 in a hypoxic incubator (Heraeus Kendro HeraCell150, Germany) for 12 hours. Cell lines were authenticated by Short tandem repeat profiling and were routinely tested to ensure the absence of human pathogens and mycoplasma contamination.

### Chemicals and reagents

Temsirolimus (S1044) and rapamycin (S1039), were purchased from Selleckchem (Houston, TX, USA). PT2385 was obtained from MedChemExpress (New Jersey, USA). Stocks were maintained at a concentration of 10 mM in sterile DMSO. DMSO served as vehicle in all experiments. Plasmids GIPZ lentiviral human PCDHGC3 shRNA (V3LHS_320145, V3LHS_320144 and V3LHS_353056) and GIPZ non-silencing lentiviral shRNA control were purchased from Horizon Discovery (Cambridge, UK).

### Generation of stable knockdown and cell transfections

Nontargeting short hairpin RNA (shRNA) or *PCDHGC3* shRNA constructs cloned into the pGIPZ lentiviral vector were purchased from Horizon Discovery Ltd. (Cambridge, United Kingdom). For viral infection of 786-O or RCC4 cells, the regular medium was replaced with culture medium containing 5 mg/mL Polybrene and cells were kept under selection for 10 days. For *PCDHGC3* overexpression, transient transfection with a mouse derived-*PCDHGC3* GFP-vector (FL-C3-GFP) was performed using Lipofectamine LTX (Invitrogen, Thermo Fisher Scientific), following the manufacturer’s protocol, at a final plasmid DNA concentration of 10 ng/μL. The FL-C3-GFP plasmid was generated as previously described (Lobas *et al*, 2012). Empty vectors were used as controls (CT). Percentage of transfected cells was quantified by flow cytometry.

### Cell invasion in 3D-bioprinted cancer models

To analyze cell invasion, we assessed the capacity of cells to invade bioinks. Bioink composition, hydrogel preparation, and bioprinting conditions are detailed in our previous report(Herrada-Manchón *et al*, 2021). To evaluate the cell invasion capacity within the bioink, 1 x 10^5^ cells were seeded on top of each scaffold. After spontaneous spheroid formation, microscopy imaging was conducted over a 40-hour period. Quantitative analysis of the spheroids’ area and perimeter was then performed.

### Cell proliferation assays

Cell growth and viability were assessed using the MTS CellTiter 96® AQueous One Solution Cell Proliferation Assay (Promega, Madison, Wisconsin, US) after 72 hours of treatments, according to the manufacturer’s instructions. Absorbance at 490 nm was measured with a 96-well plate reader after 1 hour of incubation with MTS reagent. Real-time cell proliferation was evaluated using the iCELLigence system (ACEA Biosciences, San Diego, CA, USA). 3 x 10^3^ cells were seeded onto plates containing gold microelectrodes biosensors that measure impedance when cells adhere to the microelectrodes. The impedance increases with higher rates of cell proliferation and adhesion. Drugs were added 26 hours after cell seeding, and cell impedance was monitored every two hours for 96 hours. Data were analyzed using the RTCA analysis software (ACEA Biosciences) and represented as cell index (CI), calculated as follows: CI = (impedance at time point n - impedance in the absence of cells)/nominal impedance value. To analyze cell proliferation in a 3D environment, we used the 3D-bioprinted cell model in which cells were embedded in the bioinks and then printed as scaffolds as previously described (Herrada-Manchón *et al*, 2021). Drugs were added 24 hours post-printing. Cell growth was monitored through image analysis using microscopy. Cell counting was performed on Z-stack images (comprising 15-20 sections with a 20 μm interval between them) at randomly chosen positions within the scaffold before and after 14 days of treatment. Images were acquired with green fluorescence using a Zeiss AxioObserver Z1 microscope (Carl Zeiss, Germany), a camera (AxioCam MRm; Carl Zeiss), and Apotome (ApoTome 2; Carl Zeiss).

### Colony formation assays

Cells were seeded at low density (150 cells) in twelve-well plates. After 24 hours, cells were incubated with the drugs and cultured for 7 days. Subsequently, cells were fixed with ice-cold methanol and stained with 0.5% crystal violet. The percentages of area that is covered by colonies were quantified using a plugin for ImageJ software.

### Cell cycle analysis

For the cell cycle analysis, cells were fixed in 70% ice-cold ethanol, permeabilized with 0.1% Triton X-100/PBS and stained with DAPI at a final concentration of 1 μg/mL. Flow cytometry analysis was performed to quantify the distribution of cells in the G1, S, and G2–M phases of the cell cycle.

### Murine models

Animal studies were performed in accordance with the Declaration of Helsinki and with the guidelines and regulations of the University of Oviedo and were approved by the Research Ethics Committee of the University of Oviedo (PROAE 47/2019; PROAE 10/2020). Six-week-old female athymic nude mice (Charles River Laboratories, France) were used in all experiments. For tumor xenograft experiments, 1 x 10^6^ 786-O cells were subcutaneously inoculated into the right flank of nude mice. Tumor size was measured twice weekly, and volumes were calculated using the formula 4/3πRr2, where R represents the larger diameter and r represents the shorter diameter. To investigate tumor metastasis, two approaches were used. In the first experimental metastasis mouse model, 5 x 10^5^ 786-O cells were injected into the tail vein of mice. In the second model, 1 x 10^6^ 786-O cells were orthotopically injected into the renal subcapsular space. Tumor growth and metastases development were monitored at the Preclinical Image Laboratory of the University of Oviedo using a Computed Tomography scanner (Argus CT, Sedecal, Madrid, Spain). Images were obtained at a voltage of 45 kV and 300 μA. The acquired images were reconstructed using Horos, which is a free and open-source code software (FOSS) program that is distributed free of charge under the LGPL license at Horosproject.org and sponsored by Nimble Co LLC d/b/a Purview in Annapolis, MD, USA. For drug treatment experiments, when CT tumors reached approximately 100 mm^3^, mice were randomly assigned (n = 5 per group) to receive intraperitoneal injections of either vehicle or temsirolimus at a dosage of 1.5mg/kg/day, administered 5 days per week for 2 weeks (xenograft model) or 3 weeks (orthotopic model). In all cases, isolated organs and tumors were visualized using the In Vivo Imagen System (IVIS) for GFP tumor detection according to the equipment instructions.

### Immunohistochemistry

For histological examination, samples were fixed in 4% paraformaldehyde, embedded in paraffin, and then cut into 3-4 μm sections. Hematoxylin and eosin (H&E) staining was performed by deparaffinizing the sections in xylene, rehydrating them through a series of graded alcohols, and subsequently staining with H&E using standard protocols. For immunodetection, antigen retrieval was carried out using a high-(for HIF-2α immunostainings) or low- (for Ki67 immunohistochemistry) pH EnVision™ FLEX target retrieval solution in a Dako PT link platform (Dako Denmark A/S, Glostrup, Denmark), followed by staining with a Dako EnVision™ Flex detection system. Slides were incubated with the following primary antibodies: anti-HIF2α antibody (ab199, Abcam) at 1:50 dilution for 30 min and anti-Ki67 antibody (M7240, Dako Denmark A/S) at 1:100 dilution for 20 min. Fluorescence microscopy was preformed on a Zeiss AxioObserver Z1 microscope (Carl Zeiss, Germany) with a Plan-Apochromat 40X/1.3 (NA = 1.3, working distance = 0.21mm) or Plan-Apochromat 63X/1.4 (NA = 1.4, working distance = 0.19mm) oil lens objective, a camera (AxioCam MRm; Carl Zeiss) and an Apotome (ApoTome 2; Carl Zeiss).

### RNA extraction and real-time PCR (RT-qPCR)

Total RNA was isolated from cell lines with mirVanaTM miRNA Isolation Kit (Invitrogen, Thermo Fisher Scientific) following the manufacturer’s instructions. Subsequently, cDNA was synthesized from 100 ng of total RNA using the Maxima First Strand cDNA synthesis kit for RT-qPCR (Thermo Fisher Scientific). Taqman PCR Master Mix (Applied Biosystems, Foster City, CA, USA) was used to analyze the expression of the indicated genes. Each sample was analyzed for peptidylprolyl isomerase A (PPIA) mRNA to normalize RNA input amounts and perform relative quantification. All reactions were performed in triplicate, and relative mRNA expression was normalized against endogenous controls using 2^−ΔΔCT^.

### Western blotting

Cells at 80%–90% confluence were lysed using RIPA buffer (Sigma-Aldrich, St. Louis, Missouri, US) supplemented with protease and phosphatase inhibitors. A total of 30-40 μg of proteins from the lysates were fractionated on SDS-PAGE gels and transferred to PVDF membranes (Bio-Rad Laboratories, CA, USA). Membranes were probed with rabbit anti-HIF2α (ab243861, Abcam, Cambridge, UK) at 1:250 dilution; rabbit anti-HIF1α (NB100-449, Novus Biologicals, Minneapolis, MN, USA) at 1:500 dilution; mouse anti-pan-cytokeratin AE1/AE3 (sc-81714, Santa Cruz Biotechnology, Dallas, Texas, US) at 1:250 dilution; mouse anti-N-cadherin (ab19348, Abcam) at 1:500 dilution; rabbit anti-vimentin (ab16700, Abcam) at 1:500 dilution; rabbit anti-ZEB1 (NBPI-77178, Novus Biological) at 1:500 dilution; rabbit anti-ZEB2 (NBPI-82991, Novus Biological) at 1:500 dilution; rabbit anti-Snail2 (#9585 Cell Signaling Technology, Danvers, Massachusetts, US) at 1:1,000 dilution; rabbit anti-phospho-mTOR (Ser2448) (#2971 Cell Signaling Technology) at 1:1,000 dilution; rabbit anti-mTOR (7C10) (#2983 Cell Signaling Technology) at 1:1,000 dilution; rabbit anti-phospho-p44/42 MAPK (Erk1/2) (Thr202/Tyr204) (#9101 Cell Signaling Technology) at 1:1,000 dilution; rabbit anti-p44/42 MAPK (Erk1/2) (#9102 Cell Signaling Technology) at 1:1,000 dilution; rabbit anti-phospho-GSK-3β (Ser9) (#9336 Cell Signaling Technology) at 1:1,000 dilution; rabbit anti-phospho-AKT (Ser473) (#9271 Cell Signaling Technology) at 1:1,000 dilution; rabbit anti-pan-AKT (ab8805, Abcam) at 1:500 dilution; rabbit anti-phospho-S6 ribosomal protein (Ser240/244) (D68F8) (#5364 Cell Signaling Technology) at 1:1,000 dilution; rabbit anti-S6 ribosomal protein (5G10) (#2217 Cell Signaling Technology) at 1:1,000 dilution; rabbit anti-phospho-4E-BP1 (Thr37/46) (236B4) (#2855 Cell Signaling Technology) at 1:1,000 dilution or rabbit anti-4E-BP1 (53H11) (#9644 Cell Signaling Technology) at 1:1,000 dilution. Anti-β-actin (Sigma-Aldrich) from mouse, at 1:10,000 dilution, was used as loading control. Bound antibodies were detected with IRDye 800 CW or IRDye 700 IgG secondary antibodies (LI-COR Bioscience, Lincoln, NE, USA) at 1:10,000 dilution. Odyssey Fc Imaging System (LI-COR Bioscience) was used for image acquisition and the Image Studio software for densitometry analysis.

### Proteomic analysis

Quantitative proteomic analyses were performed at the Proteomics Service of the Institute of Health Research of the Principality of Asturias (Oviedo, Spain). Briefly, 5 replicates of protein extracts per condition were obtained and precipitated using standard procedures, and subsequently subjected to trypsin digestion. Samples were analyzed by nanoLC-MS/MS on a hybrid Q-TOF ZenoTOF 7600 mass spectrometer (Sciex), connected to an Evosep One chromatographic system (Evosep). Protein quantification was calculated by summing the peak areas of the corresponding peptides. Protein groups were statistically analyzed using amica software (https://bioapps.maxperutzlabs.ac.at/app/amica). Differential expressed proteins were identified for each comparison using linear models on the log2-transformed area-normalized protein levels by limma.

### Measurements of acetyl-CoA levels

Intracellular acetyl-CoA levels were measured using the Acetyl-Coenzyme A fluorometric assay kit (Sigma-Aldrich) according to the manufacturer’s recommendations.

### Oil Red O staining

For lipid droplet staining, cells maintained at 100% confluence for 3 days were fixed with 10% formaldehyde for 1 hour, rinsed with 60% isopropanol for 5 minutes, stained with 3 mg/mL Oil Red O (ORO) for 10 min, and then washed with water three times. For drug treatment experiments, confluent cultures were exposed to drugs for 72 hours. The quantification of the ORO-stained area was performed using ImageJ software.

### Seahorse assay

Cells were seeded in XFp cell culture miniplates (Agilent Technologies) at a density of 1.2 x 10^4^ cells/well in 786-O or 1 x 10^4^ cells/well in RCC4. Twenty-four hours later, samples were analyzed to measure Oxygen Consumption Rate OCR with the Cell Mito Stress Test using the Seahorse XF HS Mini Analyzer (Agilent Technologies, Santa Clara, CA, US) following manufacturer’s instructions. Concentration of inhibitors was as follows: Oligomycin (1.5 μM), Carbonyl Cyanite-4 (trifluoromethoxy) Phenylhydrazone (FCCP) (1 μM in 786-O and 2 μM in RCC4), and Rotenone/Antimycin A (R/AA) (0.5 μM). Data were normalized with the cell number following DAPI staining. Data analysis was performed using Agilent Seahorse Analytics.

### Methylation and RNAseq data analysis

Normalized RNAseq data from ccRCC samples were retrieved from The Cancer Genome Atlas (TCGA) data portal and analyzed with KM plotter (https://kmplot.com/analysis/) (Győrffy, 2024). For DNA methylation profiling analysis, Wanderer web tool was used (http://maplab.cat/wanderer). Wanderer provides beta values and statistical comparisons of HumanMethylation450 probes for the gene of interest across normal and tumor samples from the TCGA database, as previously reported(Díez-Villanueva *et al*, 2015). To evaluate bivariate associations across the cluster groups of *PCDH* genes, a thorough automated method for exploratory data analysis software (AutoDiscovery, Butler Scientifics, Barcelona, Spain) was conducted. The suitable statistical approach was selected based on the type of data and the distribution of the variables in each case, as assessed by AutoDiscovery. This procedure was implemented in each potential subgroup of the dataset, created based on metastasis and tumor type.

### Statistical analysis

Statistical analysis was carried out using Graphpad Prism 10 software. All data are presented as mean ± standard deviation (SD) or standard error of mean (SEM) of at least two independent experiments as indicated in the corresponding Figure Legends. Two-tailed or multiple student’s t test were used to determine the statistical significance between groups. Disease overall survival was estimated using Kaplan–Meier analysis (log-rank analysis for statistical significance). For statistical analysis in correlation studies, Pearson correlation coefficient was used. *p* values less than 0.05 were considered statistically significant (* *p* < 0.05; ** *p* < 0.01; *** *p* < 0.001; **** *p* < 0.0001).

## Acknowledgements

This research was funded by Instituto de Salud Carlos III through the project PI20/01754 (co-funded by the European Regional Development Fund/European Social Fund ‘A way to make Europe’/‘Investing in your future’) and the Principado de Asturias and European Regional Development Fund through the project IDI/2021/00079. LC thanks The Spanish Ministerio de Ciencia, Innovación y Universidades for a FPU predoctoral contract. TC and JS-JG thank Consejería de Ciencia, Innovación y Universidad (Principado de Asturias) for Severo-Ochoa predoctoral contracts. We would like to thank the Proteomics Service of ISPA, the Preclinical Imaging Unit of the Animal Facility at the University of Oviedo, the Flow Cytometry Unit of ISPA, and the Molecular Histopathology Unit of IUOPA for their invaluable support and contributions to this research. Figs 2, 3, 6 and 7 were partly generated using Servier Medical Art, provided by Servier, licensed under a Creative Commons Attribution 3.0 unported license.

## Author contributions

M-DC was responsible for conceptualization. LC, TC, LS, JS-J-G, AS-P, HH-M, M-A F, EM and MDST performed the experiments. LC and TC designed methodologies. LC, TC, LS, EM and MDST carried out formal analysis. JAW provided study materials and contributed to writing the manuscript. LC and M-DC contributed to data presentation. M-DC oversaw the research activity and wrote the manuscript. M-DC and JAW critically reviewed and edited the manuscript. M-DC and acquired funding. All authors have read and agreed to the published version of the manuscript.

## Expanded View Figure legends

**Figure EV1.**
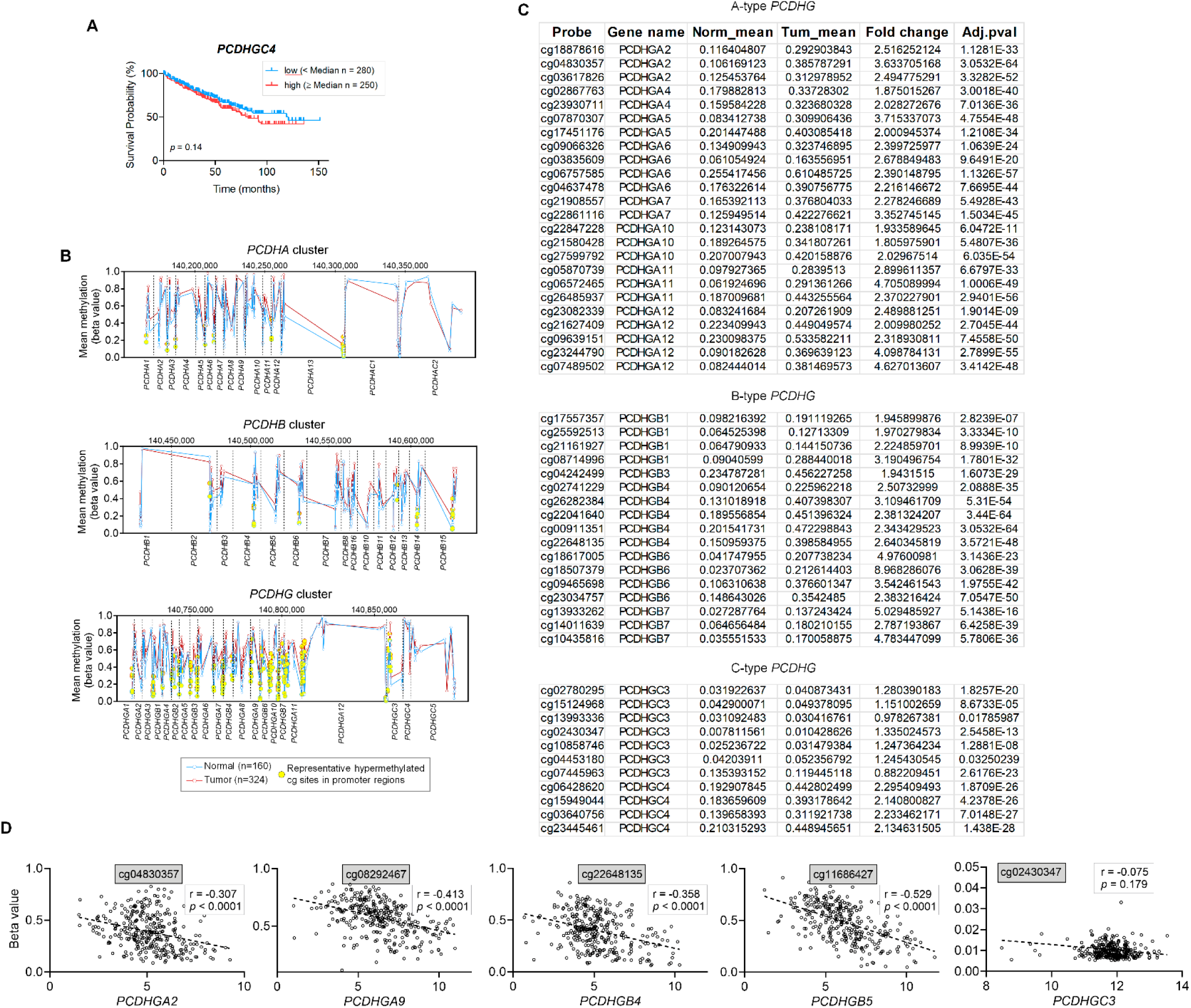
DNA methylation across the *cPCDH* genes in ccRCC included in the TCGA database. (A) Kaplan–Meier curves for ccRCC patients according to the *PCDHGC4* mRNA levels. (B) Methylation levels at CpG sites within the promoter regions of the indicated protocadherins. Median values were calculated from 324 ccRCCs (red line) and 160 non-tumoral samples (blue line). Significantly hypermethylated CpG sites are highlighted by yellow circles. Data were analyzed with the Wanderer web tool. (C) Table showing methylation levels in *g*ene promoter regions in tumor tissues and non-tumoral counterparts. (D) Correlations between methylation levels and expression of the indicated *PCDHG* genes. Pearson coefficient (r) is shown.

**Figure EV2.**
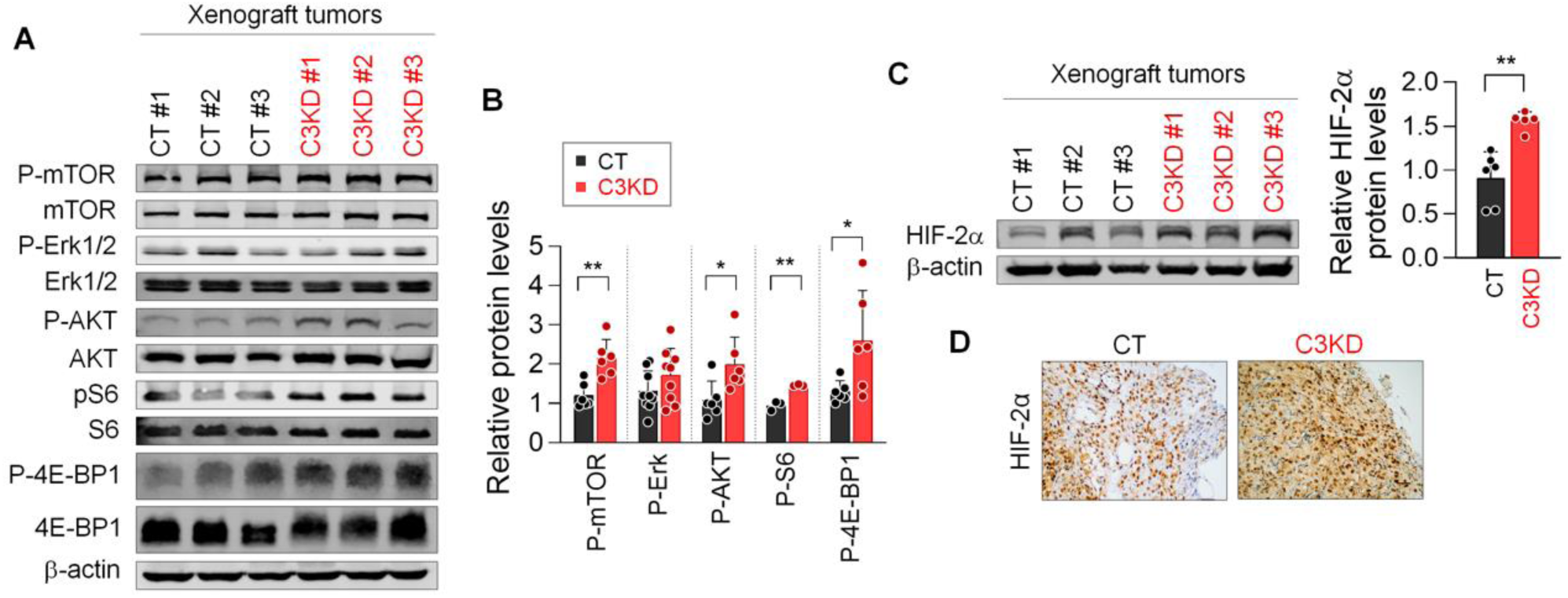
Analysis of the mTOR pathway and HIF-2α expression in tumor xenografts. (A-C) Representative images of western blots and their respective quantifications of the indicated proteins involved in the mTOR and ERK pathways (A, B) and the HIF2α protein (C) in three independent CT and C3KD derived tumor xenografts (D) Representative immunohistochemical images of HIF-2α in CT and C3KD tumor xenograft. Scale bars = 200 μm. *p < 0.05, **p < 0.01.

**Figure EV3.**
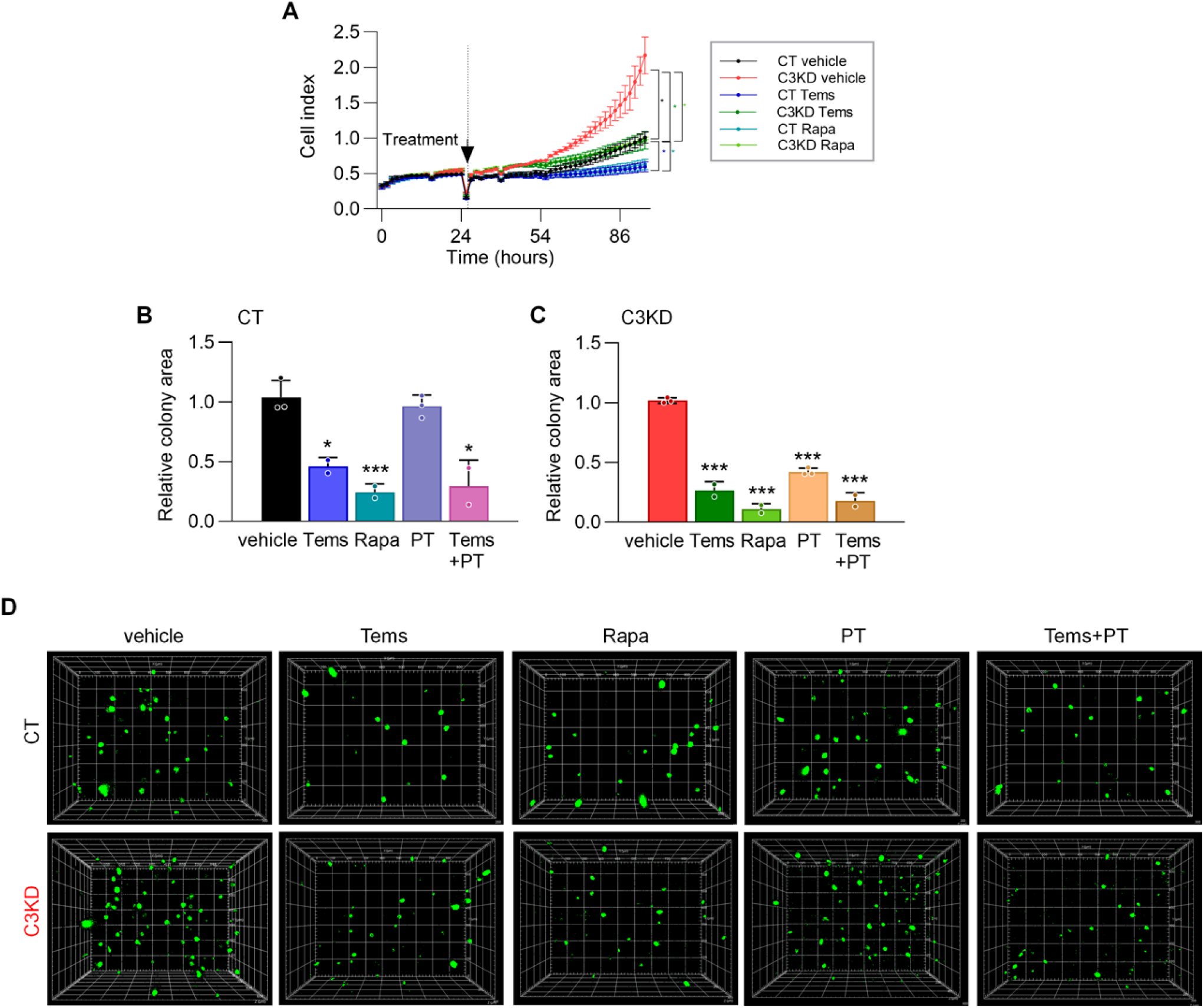
mTOR and HIF2α inhibitors suppress high proliferative rate induced by *PCDHGC3* deficiency in RCC4 cells. (A) Real-time analysis of cell proliferation with the iCELLigence system. (B, C) Colony formation assays in RCC4 CT (B) and C3KD (C) cells after treatment with the indicated drugs. (D) Representative fluorescence XYZ projections of CT or C3KD RCC4 printed cells treated with the indicated drugs. *p < 0.05, *** p < 0.001.

**Figure EV4.**
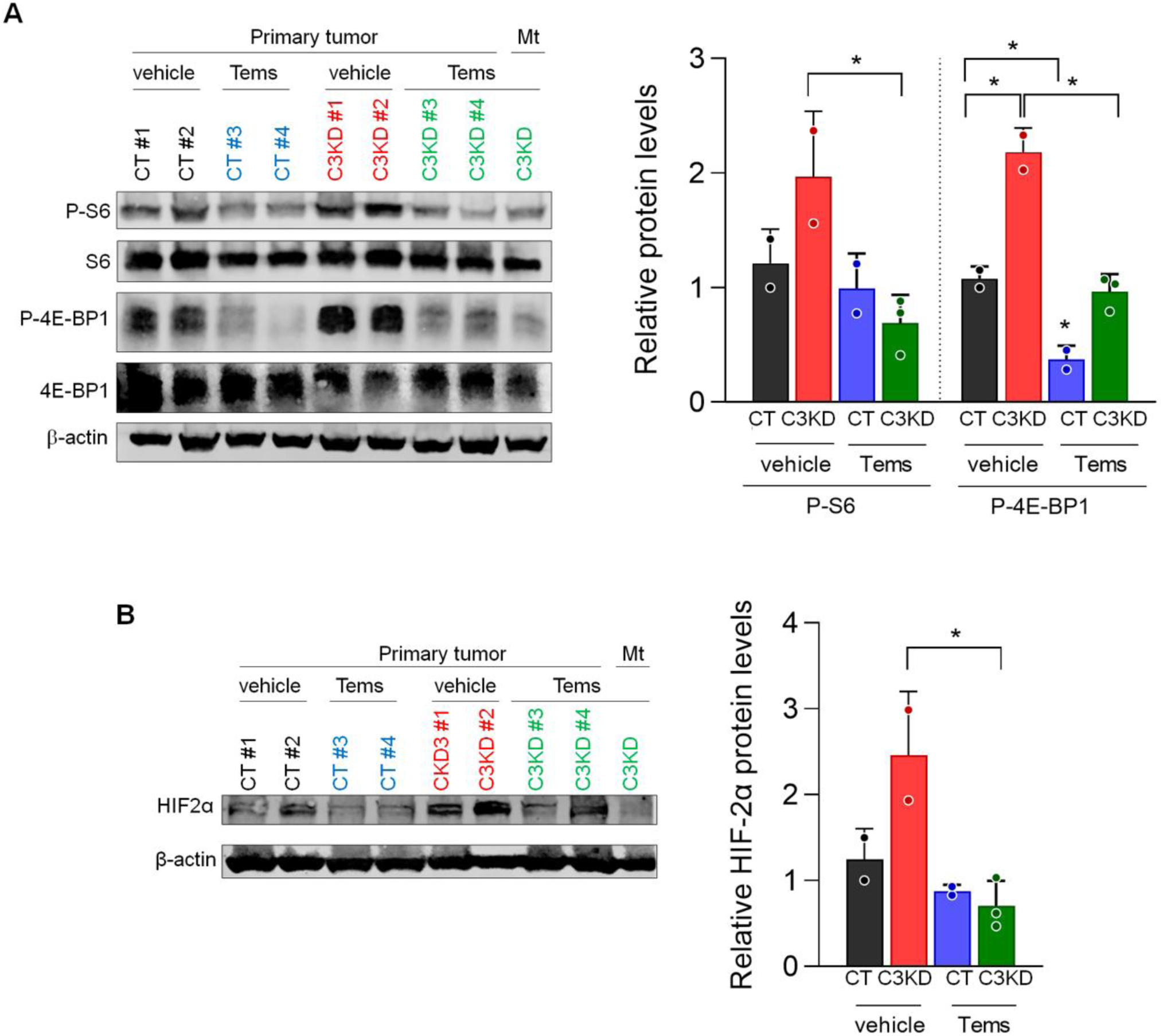
Analysis of mTOR pathway (A) and HIF-2α (B) expression in tumoral and metastatic tissues raised in the orthotopic xenograft model. (A) Representatives immunoblot images and quantifications of the indicated proteins in independent primary tumors and metastasis (Mt). *p < 0.05.

**Figure EV5.**
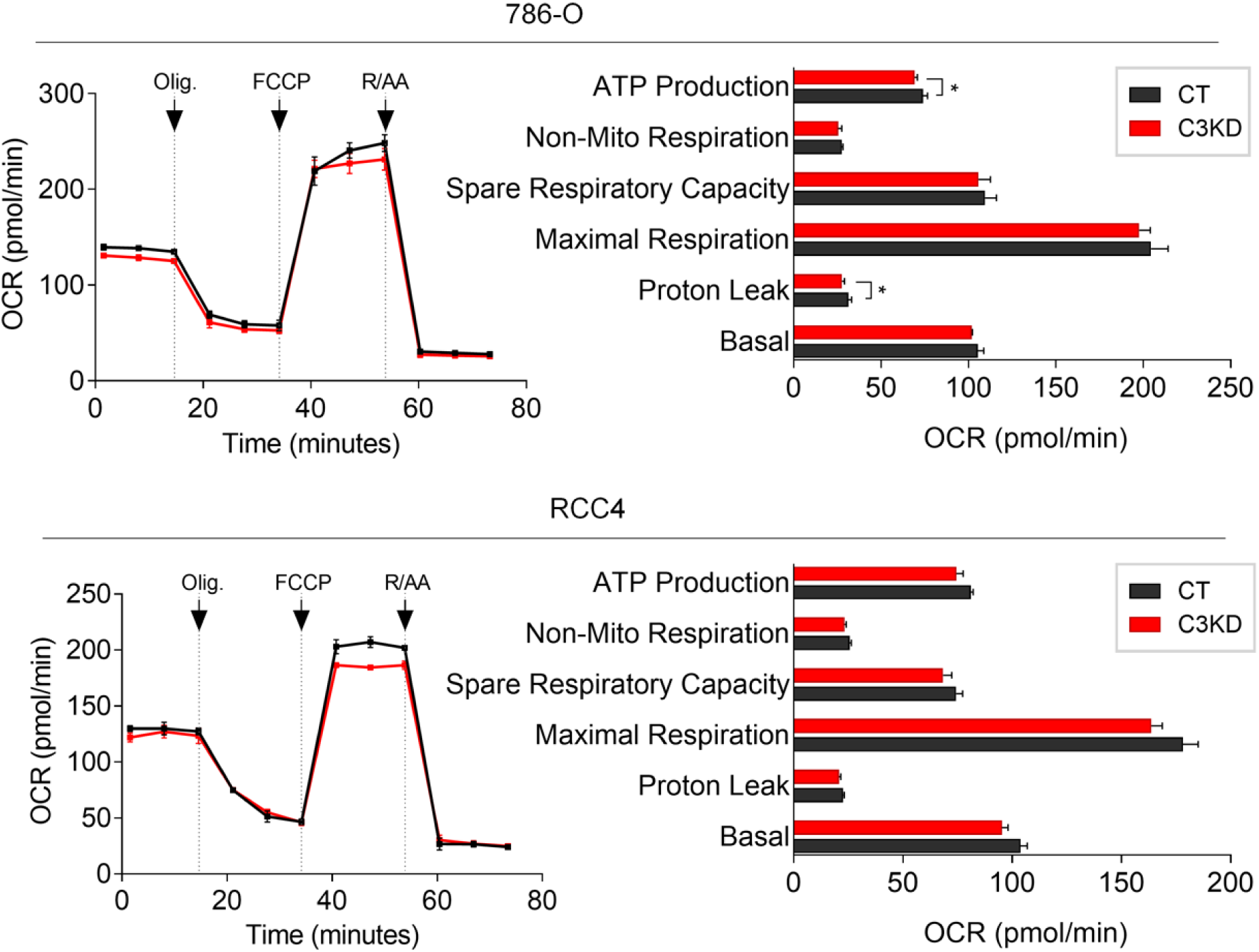
Analysis of mitochondrial function in CT and C3KD cells. Seahorse Cell Mito Stress Test assays performed in CT and C3KD 786-O and RCC4 cells. Data are represented as oxygen consumption rate (OCR). Arrows indicate additions of the indicated stressors: oligomycin (Olig), carbonyl cyanite-4 (trifluoromethoxy) phenylhydrazone (FCCP) and rotenone/antimycin A (R/AA).

**Table EV1. Proteins differentially expressed in CT versus C3KD 786-O cells.**

**Table EV2. Detailed data on GO terms and pathways enriched in CT versus C3KD 786-O cells based on proteomic analysis.**

